# The interplay of climate change, urbanization, and species traits shapes European butterfly population trends

**DOI:** 10.1101/2025.02.13.638066

**Authors:** Pau Colom, Ashley Tejeda, Simona Bonelli, Benoît Fontaine, Mikko Kuussaari, Dirk Maes, Xavier Mestdagh, Miguel L. Munguira, Martin Musche, Lars B. Pettersson, David Roy, Johannes Rüdisser, Martina Šašić, Reto Schmucki, Constanti Stefanescu, Nicolas Titeux, Josef Settele, Chris van Swaay, Javier Gordillo, Yolanda Melero

**Author notes:** Corresponding author: Pau Colom.

## Abstract

Species populations naturally fluctuate, yet long-term trend analysis can reveal patterns of success, decline, or stability under global change pressures. While responses to climate change are well-documented, its synergy with another major global driver, urbanization, remains understudied. Here, we analyzed long-term monitoring data from over 8,400 populations of 145 butterfly species across Europe, representing a high diversity of species traits, to assess population trends in response to climate change and urbanization. We examined how population responses vary between urban and rural contexts, providing insights into the influence of site-specific conditions. Climate warming was associated with population declines, which were more pronounced in urban areas. The effect of precipitation varied between environments: increases in precipitation generally benefited populations in rural areas but had detrimental effects in urban ones. Aridity consistently drove population declines across environments, with slightly stronger effects in urban areas. Species with colder climatic niches declined the most in response to warming, increased aridity, and reduced precipitation, while trophic specialists were particularly vulnerable to aridity and precipitation changes in urban environments. Although increasing urbanization did not explain overall population trends, its effects became evident when considering species traits, with certain traits being more vulnerable to urbanization. Specifically, species with narrow climatic niches declined the most in response to urbanization in rural areas, while those and larger body sizes decline the most in urban environments. Our findings highlight the complex interplay between environmental change, landscape context, and species traits in shaping biodiversity outcomes. Importantly, our results suggest that urbanization generally amplifies the impact of climate change on insect population trends.

## 1. INTRODUCTION

Urbanization and climate change are accelerating biodiversity shifts in the Anthropocene (Moorsel et al., 2022; Urban et al., 2024). In Europe, built-up surface area increased from 173.6 billion m^2^ (ca. 4% of total land area) in 1975 to 464.6 billion m^2^ (ca. 10.7%) in 2020 (EU Commission, 2023); a tendency that has expanded worldwide and is projected to increase sixfold by 2050 (Gao & O’Neill, 2020). In parallel, global temperatures have risen by 0.15 to 0.2°C per decade, and other climatic patterns (e.g. seasonality and precipitation patterns) have shifted in different directions across regions (Gulev et al., 2021).

Urbanization typically transforms natural environments into small, isolated patches, surrounded by a built matrix largely inhospitable to wildlife (Liu et al., 2016). This habitat loss and fragmentation restricts dispersal and establishment across suitable habitats, driving declines in diversity and abundance of many taxa (Dri et al., 2021; Fenoglio et al., 2020; Hou et al., 2023; Piano et al., 2020). However, some species may adapt and even thrive in urban environments by exploiting new resources or niches (i.e. urban exploiters; Korányi et al., 2021; Kurucz et al., 2021; Santana Marques et al., 2020). Likewise, climate change also disrupts the environmental conditions species are adapted to, and hence population stability (Bellard et al., 2012; Cahill et al., 2013; Parmesan, 2006; Root et al., 2003). Yet, for some species, climate change may create opportunities for expansion or increased populations in newly favorable conditions (Crossley et al., 2021; Fürst et al., 2023; Jackson et al., 2022). Understanding how urbanization and climate change influence population changes over time is critical, as the processes that affect the intrinsic population growth rates precede expansions, extinctions and changes in community composition (Collen et al., 2011). Extensive research has examined the effects of climate change on population trends, often supported by national monitoring programs and global repositories (e.g. Martay et al., 2017; Williams et al., 2022). However, fewer studies have focused on the impacts of urbanization alone or in combination with climate change. Those that do often rely on presence/absence data or opportunistic counts (but see Dennis et al., 2017; Grünwald et al., 2024), limiting our ability to accurately assess long-term population responses.

Another critical knowledge gap is understanding how the magnitude and direction of these urban- and climate-driven effects on populations vary across different contexts. The distinction between urban and rural environments highlights an important framework for understanding how site-specific conditions influence population responses. For instance, climate-driven effects may be intensified in urban environments due to the urban heat island effect and the limited availability of cold microclimates to buffer warming effects. However, rural populations are likely to be more vulnerable to increasing aridity and drought as they rely heavily on unmanaged natural resources and often lack human interventions (e.g. irrigation) that are more common in urban areas (Chen et al., 2014). Populations of the same species may also experience different impacts of increasing urbanization in rural areas compared to sites that have already undergone significant urban development (i.e. urban sites). Rural populations that are less exposed to human disturbances may experience greater instability as urbanization encroaches, whereas some species populations in urban sites may have already adapted (via plasticity or evolution) to cope with these pressures (Diamond et al., 2017; Merckx et al., 2024; Theodorou et al., 2018; Van de Schoot et al., 2024). However, even then, there may be tipping points under intensified urbanization where even well-adapted urban populations may begin to experience steep declines.

Adding further complexity, species responses to environmental drivers can vary even within the same environment type. Trait-based approaches can provide insights into the mechanisms that shape the response of species to environmental change, insights that a purely species-based approach may overlook (Zakharova et al., 2019). Species traits determine their resources use, their interactions with the environment as well as their capacity to cope with changes. For example, trophic generalist species may better cope with the reduced and shifting resources in novel and changing environments, while specialists that rely on specific resources are more vulnerable to these changes (Callaghan et al., 2020; Colom et al., 2022; Pla-Narbona et al., 2022). Dispersal capacity may also be critical: species with high mobility are predicted to navigate through fragmented habitats more easily, whereas less mobile species are often confined to isolated patches being more affected by habitat loss (Bommarco et al., 2010; Niebuhr et al., 2015; Öckinger et al., 2010). Traits related to the species climatic niche also play a crucial role in shaping species responses to environmental change. Species with a climatic niche center in warmer regions are generally more likely to withstand rising temperatures, while those with niche centers in cooler climates face greater vulnerability as conditions shift beyond their optimal range (Diamond et al., 2012; Engelhardt et al., 2022; Shirey et al., 2024). Likewise, species with broader climatic niche breadths, able to thrive across a wider range of climatic conditions, are often more resilient to fluctuations and extremes. In contrast, species with narrower climatic niches (either warm or cold centered), finely tuned to specific environmental conditions, may struggle to adapt to rapidly changing climates (Gregory et al., 2009; Herrera et al., 2018). Lastly, species with longer active periods or multiple reproductive cycles per year experience a wider range of climatic conditions, which may provide them with a broader environmental niche (Callaghan et al., 2021; Franzén et al., 2020). Ultimately, species whose traits confer greater flexibility are better positioned to succeed in changing environments, such as those brought on by urbanization and climate change, providing them with a competitive advantage in an increasingly unpredictable world (Hahs et al., 2023; Sol et al., 2024).

In this study, we investigate how species populations respond to the increase of urbanization and accelerating climate change. Further, we examine differences in rural versus urban environments, anticipating that these settings may differ in ways (e.g. landscape structure or microclimates) that influence the effects of environmental change on population trends. To achieve these objectives, we analyzed long-term count data from Butterfly Monitoring Schemes across Europe (Sevilleja et al., 2020). Butterflies are effective bioindicators for assessing the impacts of urbanization and climate shifts, not only due to their sensitivity to environmental changes but also because they are well-studied and extensively monitored, providing a wealth of high-quality data (Parmesan, 2003; Thomas, 2005). Moreover, the comprehensive understanding of the diverse life-history traits and climatic distribution of European species making them particularly useful in trait-based modelling (Middleton-Welling et al., 2020; Schweiger et al., 2014; Shirey et al., 2022). Specifically, we aim to answer the following key questions: (1) How do butterfly populations in urban and rural environments respond over time to urbanization and climate trends? (2) To what extent do species traits like trophic specialization, dispersal capacity, reproductive rate and climatic niche center and breadth shape their responses to these environmental changes?

We hypothesize that populations in rural environments will be more affected by urbanization, as potential adaptations over time will not have yet occurred. In contrast, a lack of cold microclimates in urban areas could amplify the impact of climate warming on urban populations. Urbanization reduces groundwater recharge, limits infiltration, and increases evaporation by decreasing vegetation and adding impervious surfaces. Consequently, populations in urban areas may be more vulnerable to changes in precipitation and aridity. We also expect population responses to be mediated by species traits. Specifically, we expect species with more flexible dietary preferences, higher reproductive rates, and increased dispersal capacity will be less affected by urbanization since these traits may confer pre-adaptations to novel environments, at both rural and urban environments; and species from warmer climates and/or with broader niche breadths will be less impacted by climatic pressures, as these traits are likely linked to the species capacity to tolerate higher temperatures and environmental variability driven by climate change.

## 2. METHODS

### 2.1 Butterfly population trends

We used the European Butterfly Monitoring Scheme (eBMS v5.0) dataset, which includes count data of 19 countries, leveraging the collaborative work of 21 citizen science monitoring projects (Roy et al., 2020). This version of the eBMS dataset spans from 1976 to 2021 (varying depending on the project, with a mean of 16 ± 11.3 years of data), covering 12,033 sites and 318 butterfly species across Europe (Fig. S1a). Each monitoring scheme contributing to the eBMS database involves a network of sites where professional and skilled volunteers conduct regular (mostly weekly) butterfly counts along fixed transects, following the standardized ‘Pollard Walk’ protocol (Pollard & Yates, 1993). Along these transects, all butterfly species are monitored during the butterfly flight season, which varies by climatic region from March to October.

To calculate annual abundance estimates for species populations, we used the regional GAM approach (Dennis et al., 2013; Schmucki et al., 2016). This two-stage method first fits a generalized additive model (GAM) with a Poisson distribution and log link function, capturing the overall variation in counts of butterfly species over time within a specific region and year (i.e. the flight curve). In the second stage, the standardized flight curve is used as an offset in a log-linear model to predict the value of missing counts per site and year. This allows us to generate complete series of weekly counts and derive local annual abundance indices for each species-site-year combination, using both observed and imputed weekly counts.

Regions were defined accounting for climate and photoperiod, both recognized as ecological determinants of butterfly phenology (e.g. Hodgson et al., 2011). First, we classified all sites into ten bioclimatic zones according to Metzger et al (2013). Second, we defined six latitudinal zones from 28.074N to 65.204N, ensuring a maximum day length variation of approximately 1.5 hours within regions, using the summer solstice (June 21st) as the reference day (Fig. S1a). Each region was thus defined as a unique combination of bioclimatic and latitudinal zones.

Regional GAMs were performed using the “flight_curve” function from the rbms package (Schmucki et al., 2022) for each species-region-year combination with five or more sites that met the following criteria: a minimum of ten weekly counts and the focal species observed in at least three different weeks. The maximum number of sites included to fit a regional GAM was 300 to balance computational demands and maintain a consistent and representative dataset. For cases with more than 300 available sites, we used an algorithm to select the “best informed” 300 sites based on the monitoring effort and each species occurrence, and to ensure a balanced spatial distribution of the selected sites across the region. For all species populations, we estimated and imputed the missing counts daily over the monitoring season using the corresponding regional flight curve with the function “impute_count” (Schmucki et al., 2022). We calculated annual abundance indices of species populations as the total of real and imputed counts per site and year. Finally, population trends for each species-site combination with ten or more years meeting the GAM inclusion criteria (i.e. ten weekly counts and the focal species observed in at least three different weeks) were calculated as the beta coefficient of year on the logarithm of annual abundance using generalized least squares models (GLS). An autoregressive correlation structure of order 1 was applied to the residuals to account for temporal autocorrelation. The resulting dataset included 8,409 population trends for 145 species across 869 sites in 12 different countries, covering six European climate zones (Fig. S1b).

### 2.2 Ecological and life-history species traits

We selected five traits known to predict butterfly population trends (Curtis et al., 2015; Eskildsen et al., 2015; Melero et al., 2016) and species urban affinity (Callaghan et al., 2021; Pla-Narbona et al., 2022). One trait related to trophic specialization: (i) the host-plant specialization index (HSI), which quantifies the trophic specialization of butterfly species in the larval stage based on the number of plant families, genera, and species they use. One trait related to body size (which in butterflies is often correlated with dispersal capacity; e.g. Sekar, 2012): (ii) the wing index (WI), derived from forewing length and wingspan of both females and males. Two traits related to climatic niche: (iii) the species temperature index (STI), i.e. the mean temperature within the species’ range, proxy of climatic niche center; and (iv) the species temperature variation index (STVI), i.e. the standard deviation of the temperature within the species range, proxy for the species climatic niche breadth. Finally, regarding reproductive rate we used: (v) the flight months average (FMA), measured as the average number of months of the year a species is observed in the adult stage (i.e. proxy of flight period length). Traits related to trophic specialization, body size and reproduction were extracted from (Middleton-Welling et al. (2020) and traits related to thermal tolerance from Schweiger et al. (2014). Out of the 145 species in our dataset, 19 lack data on thermal tolerance traits and two lack data on HSI (Table S1).

### 2.3 Urbanization data

To assess urbanization and its trends, we used the Global Human Settlement Layer (European Commission, 2023). For each site, we extracted the built-up surface (m^2^) from the GHS-BUILT-S R2023A raster dataset, derived from Sentinel-2 composite and Landsat data, in grids of 2x2 km with a resolution of 100x100 m pixels. To calculate urbanization trends, we first predicted the yearly built-up surface at each site by modelling the built data available in five-year intervals from 1975 to 2025, using four different models (linear, polynomial, exponential, and logarithmic). We then selected the model with the lowest AIC, with 99.3% of cases being exponential and 0.7% linear (cases where built-up surface remained zero over all the time series). Next, using the predicted data from the selected models, we calculated temporal trends for the specific subset of years corresponding to each species-site temporal series by determining the slope of the predicted values. This approach allows us to estimate the rate of change in urbanization, accurately reflecting exponential growth patterns in the data.

Using the GHS-SMOD - R2023A dataset, sites were classified as urban if they belonged to urban clusters, defined as contiguous 1x1 km grid cells (connected by any edge or corner) with a density of at least 300 inhabitants per km² of permanent land and a total population of at least 5,000 inhabitants in the cluster (European Commission, 2023). Sites were classified as rural if they were present in grid cells that did not belong to urban clusters. Most rural sites have a population density below 300 inhabitants per km²; however, some may have higher densities but do not form clusters with a sufficient total population to be considered urban. The assigned category (rural: n = 745 sites, 145 species; urban: n = 115 sites, 100 species; Fig. S1c; Table S1) for each species-site was based on the first year of the temporal series, which could vary between species at the same site if they had different starting years of their temporal series. Despite the species-site specific categorization, no site was classified as both urban and rural for different species. See Fig. S2 for urbanization trends for rural and urban sites.

### 2.4 Climate data

We used the ClimateDT tool (Marchi et al., 2024), which employs dynamic lapse-rate calculations for the spatial downscaling of climatic surfaces, to acquire highly reliable climatic data for transect sites. This tool leverages CHELSA v2.1 (Karger et al., 2017), a global dataset of climatic measurements at 30 arcsec spatial resolution (1 km up to 500 m). Its algorithm accounts for variations in temperature with elevation and incorporates orographic effects on cloud cover and radiation, making it a highly reliable source for climate estimations in topographically diverse areas. This ensures that our climate analyses are both more accurate and relevant to the specific conditions of our sites.

For each site and year, we extracted annual estimates of mean temperature, precipitation, and aridity (calculated as the inverse of the De Martonne index). These three climatic variables are important for butterfly populations as they directly affect their life cycles, habitat suitability, and food availability (Hill et al., 2021; Wilson & Fox, 2021). Further, the strength of their effect can vary depending on the species and the population (Colom et al., 2021; Melero et al., 2022; Mills et al., 2017). We then used linear models to calculate climatic trends associated with each species-site temporal series.

### 2.5 Statistical analysis

Two different sets of linear mixed models were conducted to analyze the interaction effects on butterfly population trends. The first set of models analyzed the interactions of urbanization and climate trends (separately) with site type (rural or urban), including random slopes for species to account for inter-specific variation in population responses. The second set of models analyzed the interactions of urbanization and climate trends with species traits. In all models, population trend (beta coefficient of the annual abundance models) was the response variable, and species, site and climate region were included as random factors. Additionally, we incorporated the inverse of the variance of the population trends as weights in the models to account for the varying reliability of the population trend estimates. First, we fitted four models in which we tested interactions of urbanization and climate trends (temperature, precipitation and aridity) with site type (rural vs. urban). We restricted these analyses to the 100 commonest species in both rural and urban sites to ensure comparability by avoiding biases due to different species composition in urban and rural sites (but see Fig. S3 for plot effects including all the species). Second, we built 40 linear mixed models, one for each combination of site type, urbanization and climate trend variables, and species trait (HSI, WI, FMA, STI, STVI). We conducted the models separately for rural and urban sites, using all species with available data for all five traits (rural = 126; urban = 90) to ensure comparable results between different site types and trait models. In this second set of models, we standardized all variables to a mean of 0 and a standard deviation of 1 to compare the interaction effect sizes between each urbanization/climate trend and the different traits. Quadratic terms were not included in the models, as exploratory analyses did not reveal significant nonlinear patterns between population trends and urbanization or climate shifts for most species, allowing us to maintain more parsimonious models. All models were fitted using the glmmTMB package (Brooks et al., 2017) in R version 4.3.3 (R Core Team, 2024).

We also replicated the first set of models (i.e. interactions of environmental trends with urban environment type) using phylogenetic generalized least squares (PGLS) to account for butterfly species that shared evolutionary history, which may cause similar responses to environmental changes. These PGLS models produced very similar results to the GLMMs when we excluded the variance weights for population trends. However, due to high computational demands and the inability to incorporate these weights in the PGLS models, we opted to use GLMMs for hypothesis testing and predictions.

We kept all models (the four models testing interactions between urbanization/climate trends and site type, and the 40 models testing interactions between urbanization/climate trends and species traits) and used the AIC to compare the sets of models based on its accuracy to explain population trends.

## 3. RESULTS

### 3.1 Urbanization and climate effects on rural and urban populations

Contrary to our expectations, the effect of urbanization on population trends was similar and close to stable (λ ∼ 0) or had high uncertainty (CI > β) for both rural and urban populations (Fig. 1a; Table 1; Table S2). Additionally, when we ran the models separately for each site type using all available species, we found that urbanization had no significant effect on either rural or urban populations (Table S3).

**Figure 1.**
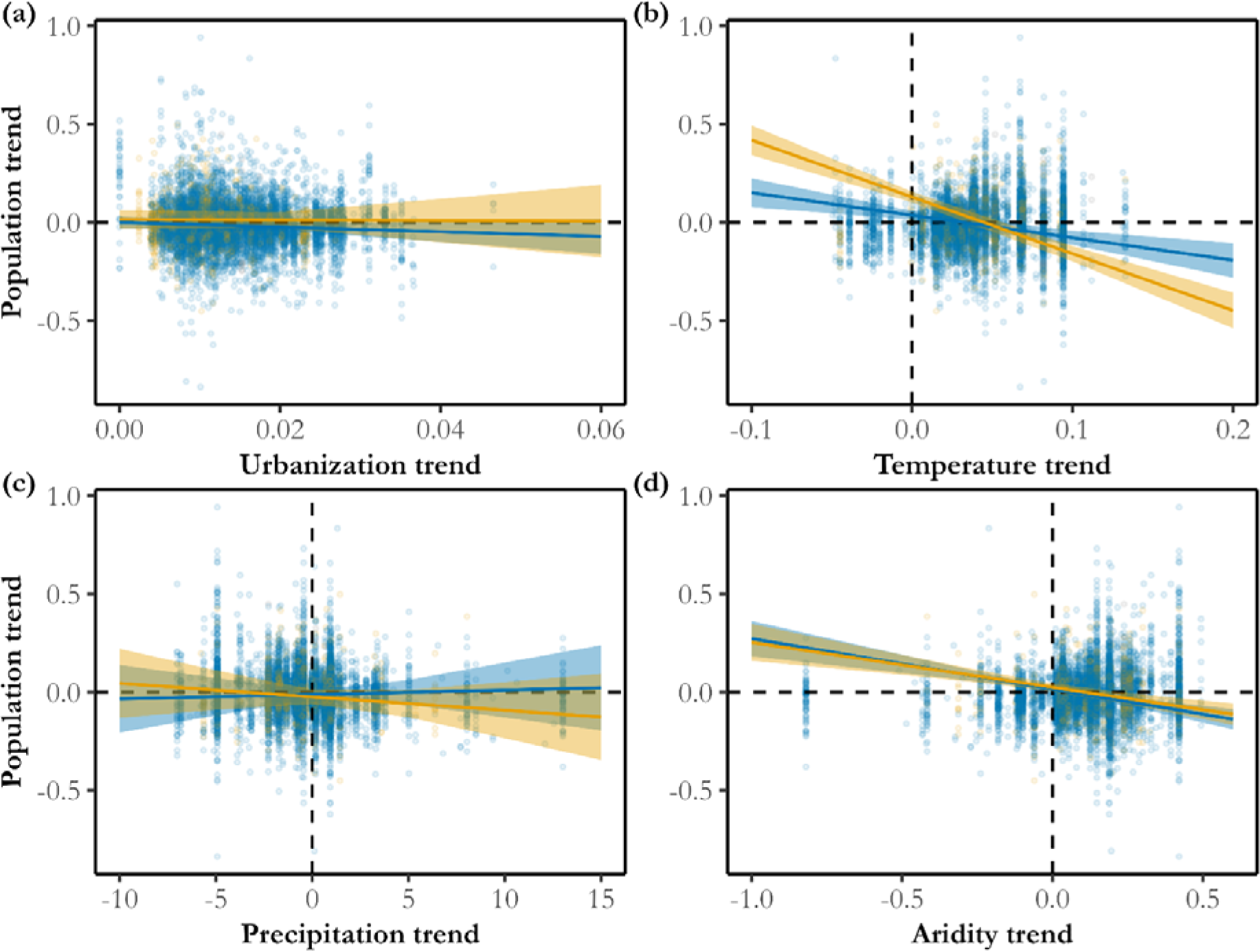
Relationships between butterfly population trends and urbanization and climate trends across rural (green) and urban (red) sites. Points represent raw data, while lines indicate model predictions with 95% confidence intervals. The models tested the interactions between each predictor: (a) urbanization trend; (b) temperature trend; (c) precipitation trend; (d) aridity trend; and environmental type (rural *vs* urban). Analyses were restricted to the 100 common species found in both environmental types across six climate regions. N = 7796 population trends.

**Table 1.** Results of interactions between (a) environmental trends (urbanization, temperature, precipitation and aridity) and environment type (rural vs. urban); and between environmental trends and species traits (HSI – host plant specialization index; WI – wing index; FMA – flight period length; STI – species temperature index; STVI – species temperature variation index) in (b) rural sites and (c) urban sites. Test statistics for each model term are shown: difference in Akaike’s information criterion AIC (ΔAIC), marginal R^2^, coefficient estimate (effect size), standard error (s.e.), z-value (Z), p-value (*P*).

| Independent variable | Interacting covariable | (a) Species in both rural and urban environments (100 species) |  |  |  | (b) Species in rural environments (126 species) |  |  |  | (c) Species in urban environments (90 species) |  |  |  |
| --- | --- | --- | --- | --- | --- | --- | --- | --- | --- | --- | --- | --- | --- |
| | | ΔAIC | $R^2_m$ | Effect size (s.e.) | Z(P) | ΔAIC | $R^2_m$ | Effect size (s.e.) | Z(P) | ΔAIC | $R^2_m$ | Effect size (s.e.) | Z(P) |
| Urbanization | Site type | 15026 | 0.007 | 1.05(2.13) | 0.49(0.62) |  |  |  |  |  |  |  |  |
| Temperature | Site type | 0 | 0.06 | -1.73(0.06) | -27.5(<0.01) |  |  |  |  |  |  |  |  |
| Precipitation | Site type | 5983 | 0.001 | -0.01(<0.01) | -15.5(<0.01) |  |  |  |  |  |  |  |  |
| Aridity | Site type | 21059 | 0.052 | 0.03(0.01) | 2.5(0.011) |  |  |  |  |  |  |  |  |
| Urbanization | HSI |  |  |  |  | 1518 | 0.001 | 0.0004(0.0003) | 13.8(<0.01) | 2813 | 0.006 | -0.003(<0.01) | 34.3(<0.01) |
|  | WI |  |  |  |  | 1498 | 0.001 | 0.0004(0.0003) | 14.4(<0.01) | 0 | 0.002 | -0.0051 (<0.01) | -63.3(<0.01) |
|  | FMA |  |  |  |  | 639 | 0.011 | 0.0009(<0.0001) | 32.5(<0.01) | 1678 | 0.003 | 0.0045(<0.01) | 48.1(<0.01) |
|  | STI |  |  |  |  | 1583 | 0.002 | 0.0003(<0.0001) | -11.1(<0.01) | 3973 | 0.003 | -0.0003(<0.01) | -3.65(<0.01) |
|  | STVI |  |  |  |  | 0 | 0.001 | 0.0018(<0.001) | 41.3(<0.01) | 3304 | 0.004 | 0.0024(<0.01) | 26.1(<0.01) |
| Temperature | HSI |  |  |  |  | 653 | 0.057 | -0.0018(<0.0001) | -39.9(<0.01) | 1489 | 0.157 | 0.0009(0.0001) | 8.1(<0.01) |
|  | WI |  |  |  |  | 1510 | 0.058 | 0.0011(<0.0001) | 27.1(<0.01) | 1554 | 0.154 | -0.0002(<0.0001) | -1.5(0.13) |
|  | FMA |  |  |  |  | 1142 | 0.06 | 0.0014(<0.0001) | 33(<0.01) | 1353 | 0.15 | 0.0017(<0.0001) | 14.2(<0.01) |
|  | STI |  |  |  |  | 0 | 0.054 | 0.0023(0.0001) | 47.4(<0.01) | 0 | 0.142 | 0.0051(<0.0001) | 39.5(<0.01) |
|  | STVI |  |  |  |  | 895 | 0.051 | 0.0015(<0.0001) | 36.8(<0.01) | 1265 | 0.165 | -0.0023(<0.0001) | -17.1(<0.01) |
| Precipitation | HSI |  |  |  |  | 854 | 0.043 | -0.0035(<0.0001) | -82.7(<0.01) | 0 | 0.015 | 0.0090(<0.0001) | 86.2(<0.01) |
|  | WI |  |  |  |  | 7595 | 0.048 | -0.0004(<0.0001) | -10(<0.01) | 7309 | 0.007 | -0.0009(<0.0001) | -8.6(<0.01) |
|  | FMA |  |  |  |  | 6531 | 0.055 | -0.0005(<0.0001) | -12.3(<0.01) | 5442 | 0.007 | 0.0045(<0.0001) | 44.1(<0.01) |
|  | STI |  |  |  |  | 0 | 0.051 | -0.0039(<0.0001) | -87.8(<0.01) | 7381 | 0.008 | -0.0001(<0.0001) | -0.8 (0.44) |
|  | STVI |  |  |  |  | 2032 | 0.05 | 0.0031(<0.0000) | 75.3(<0.01) | 6187 | 0.012 | 0.0041(<0.0001) | 34.6(<0.01) |
| Aridity | HSI |  |  |  |  | 7866 | 0.076 | 0.0022(<0.0000) | 49.8(<0.01) | 0 | 0.066 | -0.0075(<0.0001) | -67.1(<0.01) |
|  | WI |  |  |  |  | 9636 | 0.082 | 0.0011(<0.0000) | 26.6(<0.01) | 4395 | 0.078 | 0.0010(<0.0001) | 9.39(<0.01) |
|  | FMA |  |  |  |  | 9421 | 0.086 | 0.0014(0.0001) | 30.2(<0.01) | 3923 | 0.067 | -0.0024(<0.0001) | -23.7(<0.01) |
|  | STI |  |  |  |  | 0 | 0.083 | 0.0049(0.0001) | 101.8(<0.01) | 4184 | 0.074 | 0.0019(<0.0001) | 17.3(<0.01) |
|  | STVI |  |  |  |  | 8280 | 0.082 | -0.00 (<0.0000) | -45.4(<0.01) | 3407 | 0.085 | -0.0039(<0.0001) | -32.8(<0.01) |

Regarding climate, cooling trends increased population abundance over time while warming trends led to declines at both urban and rural sites. However, the negative impact of warming trends was stronger in urban than in rural populations (Fig. 1b; Table 1; Table S2). According to our model, a temperature increase of 0.02°C per year was associated with an average decrease in butterfly abundance of 2.3% per year for rural populations and of 5.7% per year in urban populations, meaning that, over one decade of climate warming, urban populations would decline by 23.4% more than rural populations.

Precipitation had opposite effects on rural and urban populations (Fig. 1c; Table 1; Table S2); for example, an increase of 10 mm/year in precipitation was associated with an average annual increase of 2.3% in rural populations, and a decrease of 6.6% per year in urban ones. Both rural and urban populations declined with increasing aridity, with rural populations showing significant steeper declines (Table 1; Table S2). However, the magnitude of the difference was relatively low (Figure 1d), for example, an increase of 0.1°C/mm was associated with an annual decline of 2.5% in rural populations and 2.2% in urban ones.

### 3.2 The role of species traits to explain urbanization and climate effects

#### 3.2.1 Inter-specific responses to urbanization trends

Although urbanization alone had no effect on rural populations (see section 3.1), it did when interacting with species traits (Fig. 2a; Table 1; Table S4). Rural population responses to urbanization were best explained by species climatic niche breadth, with overall negative responses to urbanization, but higher for species with narrow climatic niches. For instance, an increase of 1% in built-up area assuming an initial urbanized area of 100m², i.e. 2.5% of the total urbanized area (hereafter, urbanization increase), resulted in 0.09% and 0.53% annual decline of species with broader and narrower climatic niches, respectively. The interacting role with other traits was significant but almost negligible (Table 1; β < 0.001; e.g. 10 times lower than the interaction with species climatic niche breadth), with rural population overall decreasing with the increase of urbanization, yet steelier for host-plant specialists, larger, longer flight periods with climatic niche centers towards colder regions.

**Figure 2.**
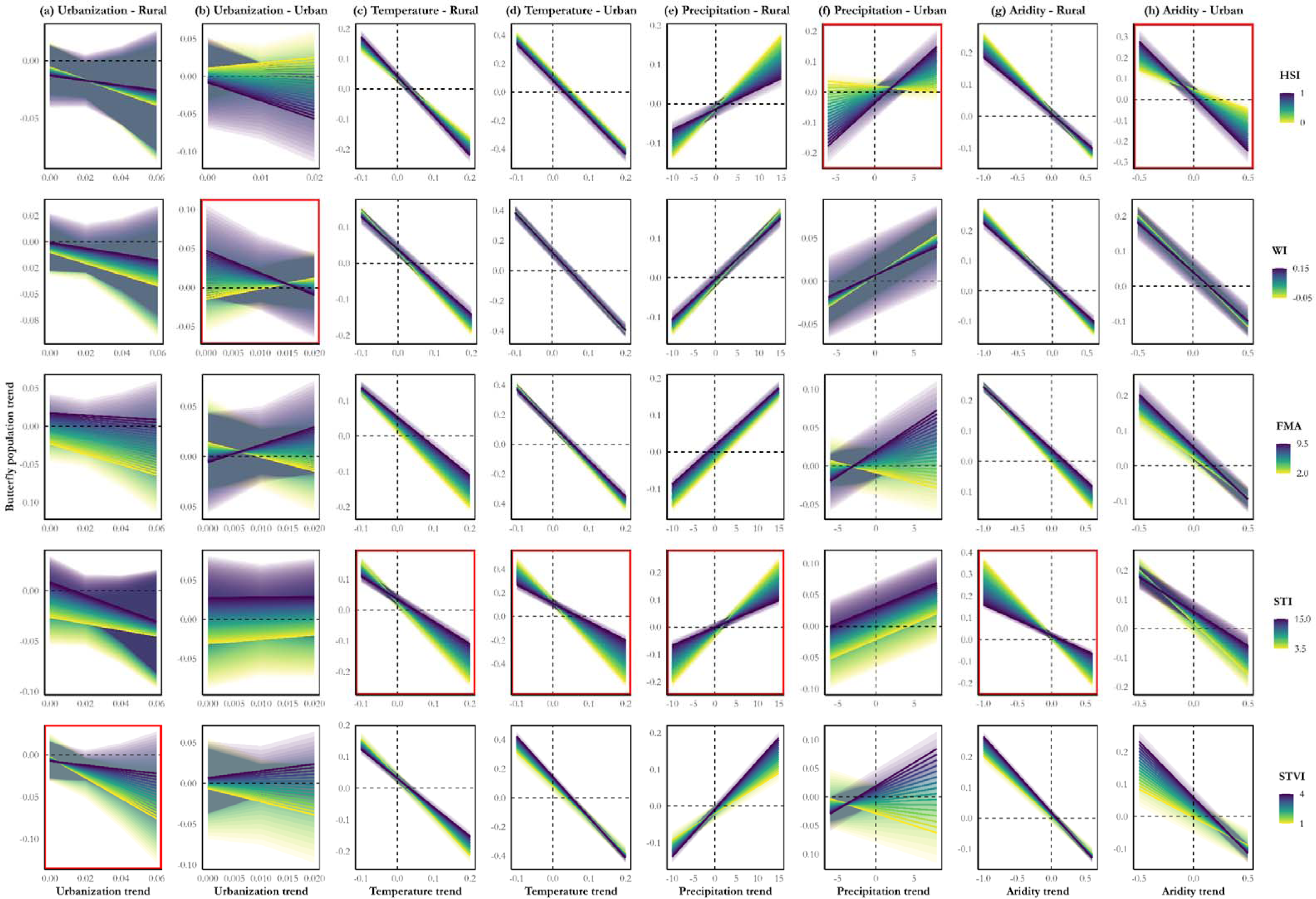
Relationships between butterfly population trends (Y-axis) and environmental trends (X-axis) interacting with species traits. Each row corresponds to a specific species trait: HSI (host-plant specialization index), WI (wing index), FMA (flight month average), STI (species temperature index), and SVI (species temperature variation index). Each column pair represents a unique predictor (urbanization (a, b), temperature (c, d), precipitation (e, f), aridity (g, h)) for rural and urban sites separated in different models. Lines represent model predictions for 20 intervals of the species trait range, with shaded areas representing 95% confidence intervals. Dashed lines indicate a trend value of 0, separating positive from negative trends (e.g., warming *vs*. cooling for temperature trends in X-axis). For each set of models (each column) the top-ranked models based on AIC are highlighted in red. Models included only species with available data for all five species traits (rural = 126; urban = 90).

Species traits in urban sites also played an important role, explaining highly divergent responses to urbanization (Fig 2b; Table 1; Table S4). Body size was the trait best associated with the response of urban populations to urbanization. Large species declined 0.44% annually, while small species increased by 0.57% with urbanization increase. Species with long flight periods were positively affected by urbanization, shifting to negative with the reduction of the flight period; as such, urbanization increased 0.82% and decreased 0.45% annually the urban populations of species with longest and shortest flight periods, respectively. Urbanization also had contrasting effects on population trends depending on their species climatic niche breadth; increasing butterfly abundance 0.64% per year in species with broad climatic niches and declining by 0.27% in the species with narrowest climatic niches. The interacting role of host plant specialization and climatic niche center was almost negligible (β < 0.001).

#### 3.2.2 Inter-specific responses to temperature trends

Increasing temperature decreased the abundance in both rural and urban populations, with species traits mediating the strength of the declines (Fig 2c, d; Table 1; Table S4).

Species climatic niche center was the main trait interacting with warming at driving population declines in rural sites. The impact of warming intensified as the species temperature index decreased, meaning temperature increases had a stronger effect on species with climatic niches centers in cold regions (hereafter, cold-centered species). For example, a temperature increase of 0.01°C per year (hereafter, temperature increase) led to an average population decrease of 0.72% annually for the warmest-centered species, but up to 1.56% per year for the coldest-centered. Overall, the temperature increase had a greater impact on host plant specialists (1.27% vs. 0.95% annual decline in generalists), those with narrower climatic niches (1.19% vs. 0.9% in broader-niche species), shorter flight periods (1.04% vs. 0.86% in longer-flight species), and larger body sizes (1.08% vs. 0.91% in smaller ones).

The climatic niche center was also the trait that best explained the variability in species response to temperature trends in urban sites, showing a similar pattern to what was observed in rural areas; the warmest-centered species declined by 1.3% per year, and up to 3.3% for the coldest-centered species with temperature increase. However, opposite to rural sites, species with broader climatic niches declined more with temperature increase (2.8% vs. 2.2% annually). The flight period length played a similar role in urban populations as in rural ones, with annual declines of 2.4% for species with the shortest flight periods and 2.7% for those with the longest in response to temperature increase. Host plant specialization and body size, also mediated the impact of temperature trends in urban populations, but with much lower effect (β < 0.001).

#### 3.2.3 Inter-specific responses to precipitation trends

Precipitation trends had more consistent positive effects on rural populations, whereas urban populations showed more variable responses, with both positive and negative impacts depending on specific traits (Fig. 2e, f; Table 1; Table S4).

Rural population responses to precipitation trends were also best explained by the species climatic niche center, with the strongest impact on colder-centered species. For example, an increase of 10mm/year in rural sites (hereafter, precipitation increase), increased population size from 6.8% up to 18.9% for the coldest-centered species. Precipitation trends had also stronger positive effects on host plant generalists than on specialist species; with e.g. annual increases of 4.5% to 13%, respectively, with precipitation increase. Likewise, the positive effect of precipitation was stronger in species with broader versus narrower climatic niches; with annual increases ranging from 6.6% to 14.2% from narrower to wider niches with precipitation increase. Species with longer flight periods and those with larger wing size were slightly more affected by precipitation trends (β < 0.001).

Precipitation effects were highly variable among species in urban sites. Host plant specialization best explained this variation, but in the opposite direction to rural sites— host plant specialists were more impacted. Host plant generalists declined by 2.3% annually, while most specialists increased up to 26.9% per year with precipitation increase. The effect also contrasted depending on flight period length, with populations declining by 1.8% annually for species with the shortest flight periods and increasing by 7.1% for those with the longest with precipitation increase. Like rural populations, urban population responses were stronger in species with broader climatic niches, with annual increases of up to 8.6%, while species with the narrowest niches declined by 3.2% in response to precipitation increase. Both body size and climatic niche center explained some variation in population responses to precipitation trends, but with considerably smaller effects (β < 0.001).

#### 3.2.4 Inter-specific responses to aridity trends

Both rural and urban populations declined with increasing aridity, but the traits shaping the strength of these declines did not always interact with aridity in the same way in the two environments (Fig. 2g, h; Table 1; Table S4).

Similar to temperature and precipitation models, species climatic niche center was the key trait explaining the effects of aridity trends in rural sites, showing that species with colder niche centers are also the most responsive. As an example, an increase in aridity of 0.1°C/mm per year (hereafter, aridity increase) resulted in annual declines of 1.4% up to 3.4% for the warmest and coldest-centered species, respectively. Increasing aridity also had stronger impacts on host plant generalists (2.4% vs. 1.6% annual decline in specialists), species with broader climatic niches (2.5% vs. 1.8% in species with narrower niches), those with shorter flight periods (2.4% vs. 2.1% in species with shorter flight times) and those with smaller body size (2.09% vs. 2.24% in larger species).

As with the precipitation models, host plant specialization was the best trait explaining population response variation to aridity trends in urban sites. In contrast to rural populations, urban populations of host plant specialist species were more affected by aridity trends than generalists, e.g. declining by 1.9% up to 5.1% per year, as species specialization increases, with aridity increase. Similar to rural populations, aridity increase led to stronger population declines in species with broader climatic niches (3.4% vs. 1.8% in species with the narrowest niches) and those with colder-centered niches (3.6% vs. 2.4% in species with warmer niche centers). However, the flight period length in rural populations interacted in the opposite direction with aridity trends compared to rural ones, with species having longer flight periods being more affected; e.g. declining by 3% per year compared to the 2.3% annual decline in species with shorter flight periods. The role of body size mediating population responses to aridity trends in urban sites, though statistically significant (p < 0.01), was almost negligible (β < 0.001).

## 4. DISCUSSION

Our analysis of long-term butterfly population trends across six bioclimatic regions reveals significant, contrasting effects of environmental pressures in rural and urban environments. Warming had the strongest negative impact on butterfly trends, with urban populations being particularly affected. The effects of precipitation change showed greater variability, with a general tendency toward positive impacts in rural areas and negative impacts in urban areas. The combined effects of rising temperatures and reduced precipitation, reflected in increased aridity, were associated with population declines across both environments, though slightly steeper in the rural ones. Contrary to our expectations, we did not find a significant effect of urbanization on population trends in either rural or urban environments. Trait modelling helped uncover the variability in species responses to these environmental changes within rural and urban environments.

### 4.1. General effects of urbanization and climate on urban and rural populations

The stronger impact of warming on urban population supports the hypothesis that urban areas generally exacerbate thermal constraints on insects. While some species may possess pre-adaptations or evolved traits in urban environments (e.g. heat tolerance in a common European moth; Merckx et al., 2024) that may help to cope with thermal stress, most insects rely on adjusting their behavior and utilize cold microclimates to optimize thermoregulation and buffer the impact of warming (Bladon et al., 2020; Suggitt et al., 2018; Vives-Ingla et al., 2023). However, the urban heat island effect, coupled with increased habitat fragmentation, may limit access to cold microsites, making it increasingly challenging for most species to cope with intensifying climate warming (Urban et al., 2024). That said, certain pollinators—such as bees—can benefit from warmer urban microclimates (Hall et al., 2017). Overall, our results suggest that the impact of warming will escalate as urban expansion continues, imposing greater thermal stress on insect populations.

Precipitation influences butterfly populations by e.g. affecting the availability and quality of their food plants, as well as the timing of plant growth and flowering (Carnicer et al., 2019; Dalton et al., 2023; Donoso et al., 2016; van Bergen et al., 2020). In rural areas, precipitation showed on average a positive relationship with population trends (see also Fig. S3), potentially reflecting the beneficial effects of precipitation on butterflies via increasing the availability and quality of their nectar-source and host plants. However, this relationship was absent, or even negative, in urban sites. Rural populations are likely more vulnerable to changes in precipitation as they often depend on unmanaged natural resources, whereas populations in urban sites typically benefit from human interventions, such as irrigation, that help mitigate water scarcity (Chen et al., 2014). Explaining a negative effect of increasing precipitation on urban populations, however, is more challenging. One possible explanation is that an increase in the number of precipitation days may reduce the time available for essential activities like feeding and mating, potentially leading to negative impacts on their fitness. Indeed, some studies have found negative correlations between increased precipitation and population size in certain butterfly species, particularly when the precipitation increase occurs during the adult stage of the butterfly (Roy et al., 2001; Ubach et al., 2022). In rural sites, however, a stronger positive effect of precipitation on resource availability may counterbalance the negative effect on time for foraging and reproduction.

Aridity can provide a more comprehensive view of how temperature and precipitation factors interact to shape butterfly population trends. While the individual effects of temperature and precipitation differed notably between rural and urban environments, the combined effect had a more consistent impact (i.e. increasing aridity correlate population decline in both environment types), though significantly stronger in rural sites. The impact of aridity could be reduced in urban environments that feature carefully managed and irrigated gardens and other green spaces (Baldock et al., 2019).

While climate shifts had clear, directional impacts on butterfly populations, urbanization itself did not significantly influence the overall population trends either in rural or urban sites. Several studies have shown that urban growth is a major factor shaping insect species composition and population abundance across spatial gradients (Knop, 2016; Kuussaari et al., 2021; Maes et al., 2022; Merckx & Van Dyck, 2019; Pla-Narbona et al., 2022; Tzortzakaki et al., 2019). However, there has been no prior evidence of urbanization effects on decadal population trends. In this study the lack of effect of urbanization on population trends could be due to (i) the filtered urban community (Gathof et al., 2022; Pla-Narbona et al., 2022; Roper-Edwards & Hurlbert, 2024), in which several, most vulnerable, species do not appear in environments suffering from urban development; and (ii) the relatively slight increase in urbanization over our study period (Fig. S2), which may not have been sufficient to drive further shifts—especially in rural sites with very low increase in urbanized area. Moreover, urban populations and their dynamics could be the result of connected populations (e.g. metapopulations) or neutral processes (i.e. their abundances reflect those of the surrounding natural populations, via e.g. source-sink dynamics), in which cases population trends will be similar or equivalent to those of the natural communities (Grünwald et al., 2024; Sol et al., 2014, 2024). In any case, our analyses suggest that the main reason for the lack of an overall urbanization effect is the high variability in interspecific responses linked to specific species traits (see section 4.3).

### 4.2 The role of species traits to modulate population responses to climate shifts

All population responses to the above-mentioned environmental pressures were mediated by species traits, although the set of traits and their strength varied among pressure and environmental type (rural versus urban). Our results show that species climatic niche is key to predicting population responses to climate shifts. Climatic traits (species climatic niche center and breadth) can be interpreted together to differentiate the effects of environmental pressures on warm-adapted species (warm and narrow niche center), cold-adapted species (cold and narrow niche center), and climatic generalists (warm/cold and broad niche center). Under global warming, cold-adapted species are declining in abundance and contracting their distribution ranges more rapidly than warm-adapted species (Bowler et al., 2015; Hällfors et al., 2024; Shirey et al., 2024), as rising temperatures push them beyond their thermal limits (Trisos et al., 2020). Butterfly trends in rural sites clearly follow this pattern with cold-adapted species being the most affected by temperature trends. In urban sites, the impact of temperature trends also increased towards species distribution centered in cold regions, however, this effect was also larger for species with broader niche centers (i.e. climate generalists). The same pattern was found for the effect of precipitation in rural populations and for aridity in both rural and urban populations. The stronger effects observed for species with colder but also broader niche centers may partly result from a confounding effect, as these two traits are moderately correlated (r = -0.37; Fig. S4). In our European multi-species dataset, butterfly species with broader niches tend to be centered in colder regions. This correlation likely arises because climatic traits were calculated based only on the European distributions of these species (Schweiger et al., 2014). Cold-centered species have most of their climatic range within Europe, while warm-centered species—such as *Pyronia cecilia*, *Zerynthia rumina*, or *Glaucopsyche melanops*—have a significant portion of their range in Africa, which was not included in the climatic niche calculations. Consequently, this likely led to a relatively overestimation of cold-centered niche breadths compared to warm-centered species. Further, species adaptations (at the population level) to local climatic conditions occur in many butterfly species (Melero et al., 2022; Roy et al., 2015), making them especially vulnerable to climate independently of their climatic niche center and breadth (which are defined at the species level), and of their location (Melero et al. in press)

Species with shorter flight periods, which depend on narrower time windows to complete their life cycle and, therefore, are likely more sensitive to changing conditions, were more significantly impacted by climate shifts in rural sites. Butterfly species with longer flight periods, typically multivoltine species (i.e. more than one generation per year), in contrast to strictly univoltine species (i.e. only one generation per year), can increase their number of generations in response to warming (Altermatt, 2010), which, while often insufficient to reverse population declines, appears to be an adaptive mechanism to climate change (Macgregor et al., 2019; Michielini et al., 2021; Wepprich et al., 2024). In urban sites, this pattern was less consistent: we found that increasing temperatures were linked to steeper declines in species with shorter flight periods, while those with longer flight periods were more affected by changes in aridity. Meanwhile, rising precipitation positively influenced species with longer flight periods but negatively impacted those with shorter flight periods. The fact that the negative impact of precipitation is observed only in species with short flight periods (i.e. short time window for foraging and reproduction) supports our hypothesis that increased precipitation can negatively impact population size by limiting the time available for essential activities (see section 4.1).

Since precipitation and aridity are more closely linked to the quality and availability of butterfly host plants than temperature (Carnicer et al., 2019; Dilts et al., 2019), it is unsurprising that their effects vary more with species trophic specialization than temperature effects. In urban sites, where trophic specialization was the strongest predictor of species responses to precipitation and aridity shifts, trophic specialists were more affected than generalists. This is likely due to their dependence on a limited number of host plant species, making them more vulnerable to shifts in resource availability and/or quality caused by changing climate conditions. However, unexpectedly, the opposite pattern was observed in rural sites, where generalists were more sensitive to precipitation and aridity shifts.

Among the traits analyzed, body size was the weakest predictor of species responses to climate shifts. This suggests that, although dispersal capacity can be crucial for adjusting geographic ranges in response to climate change (Pöyry et al., 2009), it would be less directly related to a species ability to cope with local environmental change. Traits related to climatic niche, diet breadth, and reproductive strategy provide more direct mechanisms for species to adapt and thrive under changing environmental conditions. However, body size may still be important at the intraspecific level in shaping individual responses to environmental change. Larger individuals typically gain advantages such as enhanced dispersal ability, mating success, fecundity, and survival, positively influencing overall fitness (Beukeboom, 2018; Blanckenhorn, 2000; Honek, 1993; Kingsolver & Huey, 2008). However, these benefits may be offset by longer development times and increased risks. For example, Gotthard et al. (2007) showed that adverse weather limits oviposition in *Pararge aegeria*, favoring smaller sizes, while Stefanescu et al. (2024) found that larger *Vanessa cardui* individuals face higher predation risks. Therefore, predicting whether smaller or larger individuals will better cope with environmental pressures is challenging, as it depends on the balance of associated risks and benefits of being larger or smaller.

### 4.3 The role of species traits to modulate population responses to urbanization trends

Although urbanization did not have a significant overall effect on butterfly population trends in either rural or urban environments, population responses varied significantly in relation to species traits, highlighting considerable variability at the interspecific and likely intraspecific levels. Interestingly, our results indicate that in rural sites, warm-adapted species (i.e. species with warmer and narrower niche breadths) showed the strongest declines with increasing urbanization. This pattern was not observed in urban sites, where climatic traits (at least species climatic niche center) played a less significant role. These findings reveal an unforeseen vulnerability of warm-adapted species to urbanization, suggesting a complex interplay between climatic adaptations and urbanization that warrants further investigation. Responses in urban sites were more variable: while some species showed negative trends, others experienced positive responses. Species exhibiting positive responses were the species with smaller body sizes, broader climatic niches and longer flight periods, while those with opposite traits exhibited negative responses. In contrast to its minimal influence on climate responses (see section 4.2), body size played a more significant role shaping urbanization effects (being the most important trait shaping urbanization responses in urban sites; Fig. 2b): In urban environments, larger-bodied species experienced more pronounced declines, while smaller species appeared to benefit from urbanization. These results contrast with previous findings on shifts toward larger butterflies and moths, linked to the advantage of greater dispersal capacity in areas with low connectivity among ecological resources (Kuussaari et al., 2021; Merckx et al., 2018; Pla-Narbona et al., 2022). Our results suggest that other factors related to body size rather than dispersal capacity may explain the interspecific response of butterflies to urbanization. The result of flight period length shaping species responses to urbanization aligns with previous studies that reported a stronger urban affinity in Lepidopterans with longer flight periods (Callaghan et al., 2021; Franzén et al., 2020). Similar to the impact of climate change on ectotherms phenology, urban warming can also extend the seasonal window available for adult activity in multivoltine species (i.e. species with long flight periods) (Merckx et al., 2021), potentially leading to “demographic bonanza” (Kerr et al., 2020), though theoretically this can also lead to a developmental trap (Van Dyck et al., 2015).

## ACKNOWLEDGEMENTS

This work has been made possible by the participation of more than one hundred thousand citizen scientists involved in coordinated monitoring projects across Europe over the past four decades. The UK BMS is organized and funded by Butterfly Conservation, the Centre for Ecology and Hydrology, British Trust for Ornithology, and the Joint Nature Conservation Committee. The Irish BMS is funded by the Heritage Council and the Department of Culture, Heritage and the Gaeltacht. The Dutch BMS is a co-operation between Dutch Butterfly Conservation and Statistics Netherlands (CBS), part of the Network Ecological Monitoring (NEM) and financed by the Ministry of Agriculture, Nature and Food Quality (LNV). The German BMS is a cooperation between the Helmholtz Centre for Environmental Research - UFZ, German Butterfly Conservation (GfS) and science4you. The Catalan BMS is funded by the Department of Climate Action, Food, and Rural Agenda of the Government of Catalonia and the Barcelona Provincial Council. The Finnish BMS is coordinated by the Finnish Environment Institute and funded by the Finnish Ministry of Environment. The Swedish BMS is coordinated by Lund University and is funded by the Swedish Environmental Protection Agency. The BMS Spain is coordinated by the association SOCEME, with the collaboration of Autonomous University of Madrid (UAM), Spanish Ministry of Environment (MITECO), Spanish National Parks Agency (OAPN), Sierra Nevada Global Change Observatory, and ICTS-Doñana (Doñana Biological Station, CSIC). The ZERYNTHIA BMS is supported by the Basque Country Government, Cantabria Government and Valle de Aranguren Council (Navarre). The Luxembourg BMS is coordinated by the Luxembourg Institute of Science and Technology and financially supported by the Ministry of the Environment, Climate and Biodiversity. The study and PC were funded by the research projects MEDYCI Grant PID2020-113133RB-I00 funded by MCIN/ AEI /10.13039/501100011033, and SATURNO Grant CNS2022-135489 funded by MICIU/AEI /10.13039/501100011033 and by the European Union NextGenerationEU/PRTR.

