## Supporting information for "The interplay of climate change, urbanization, and species traits shapes European butterfly population trends"

**Running title**: Climate, urbanization & population trends

Pau Colom^1^, Ashley Tejeda^1^, Simona Bonelli^2^, Benoît Fontaine^3^, Mikko Kuussaari^4^, Dirk Maes^5,6^, Xavier Mestdagh^7^, Miguel L. Munguira^8,9^, Martin Musche^10^, Lars B. Pettersson^11^, David Roy^12^, Johannes Rüdisser^13^, Martina Šašić^14^, Reto Schmucki^12^, Constanti Stefanescu^15,16^, Nicolas Titeux^7^, Josef Settele^10^, Chris van Swaay^17^, Javier Gordillo^16^ & Yolanda Melero^1,16^

^1^Department of Evolutionary Biology, Ecology, and Environmental Sciences, University of Barcelona, Barcelona, Spain. Biodiversity Research Institute (IRBio), Barcelona, Spain

^2^Department of Life Science and Systems Biology, University of Turin, Italy

^3^Patrinat & CESCO UMR7204, MNHN-OFB-CNRS-SU, Paris, France

^4^Finnish Environment Institute (Syke), Helsinki, Finland

^5^Research Institute for Nature and Forest (INBO), Brussels, Belgium

^6^Radboud Institute for Biological and Environmental Sciences (RIBES), Nijmegen, Netherlands

^7^Luxembourg Institute of Science and Technology, Esch-sur-Alzette, Luxembourg

^8^Centro de Investigación en Biodiversidad y Cambio Global (CIBC-UAM), Madrid, Spain

^9^Departamento de Biología, Universidad Autónoma de Madrid, Spain

^10^Helmholtz Centre for Environmental Research (UFZ), Department of Conservation Biology and Social-Ecological Systems, Halle, Germany

^11^Biodiversity and Evolution, Department of Biology, Lund University, Lund, Sweden

^12^UK Centre for Ecology & Hydrology, Wallingford, Oxfordshire OX10 8BB, UK

^13^University of Innsbruck, Department of Ecology, Austria

^14^Croatian Natural History Museum, Zagreb, Croatia

^15^Natural Sciences Museum of Granollers, Granollers, Catalonia, Spain

^16^CREAF, E08193, Bellaterra, Cerdanyola del Vallès, Catalonia, Spain

^17^De Vlinderstichting - Dutch Butterfly Conservation, Wageningen, Netherlands

**SUPPLEMENTARY FIGURES**

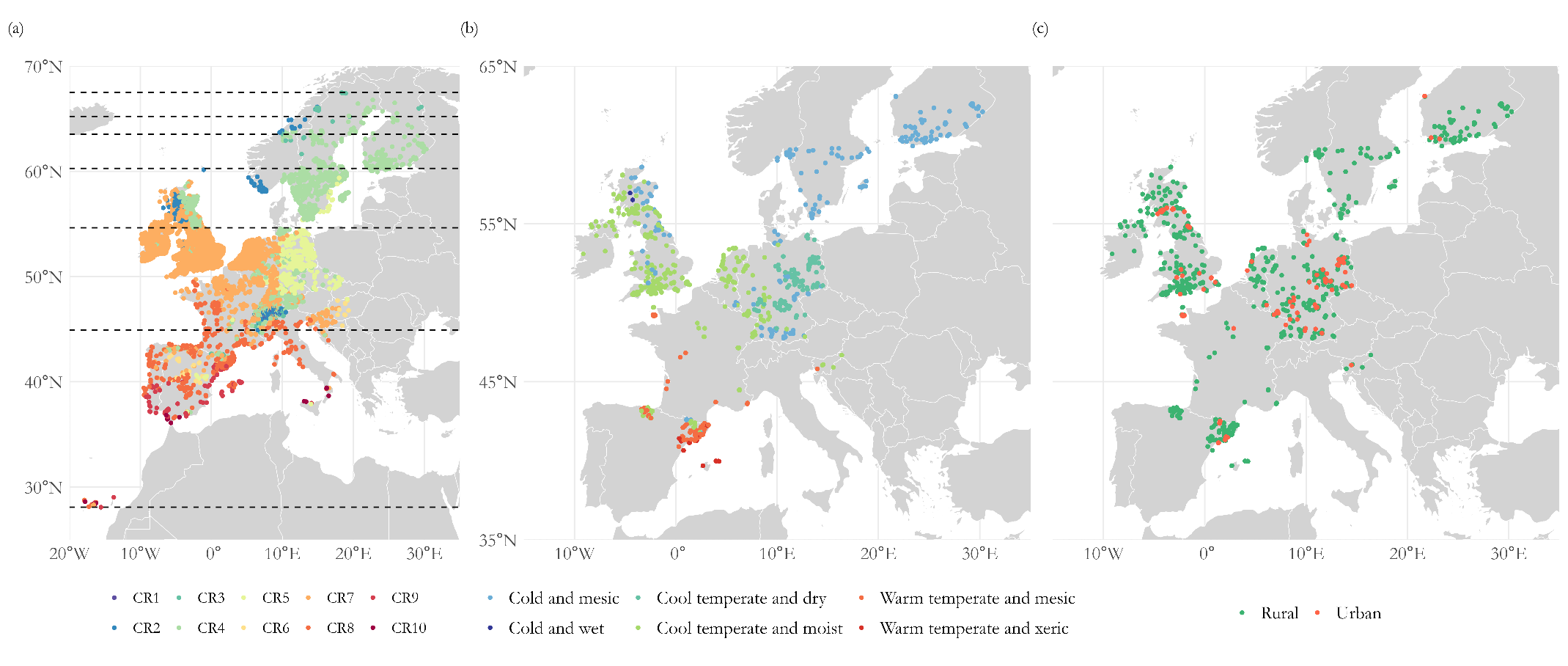

**Figure S1.** Spatial distribution of study sites across Europe categorized by climatic regions and site types. (a) eBMS sites classified into ten bioclimatic zones (Metzger et al., 2013): CR1 (extremely cold and wet), CR2 (cold and wet), CR3 (extremely cold and mesic), CR4 (cold and mesic), CR5 (cool temperate and dry), CR6 (cool temperate and xeric), CR7 (cool temperate and moist), CR8 (warm temperate and mesic), CR9 (warm temperate and xeric), and CR10 (hot and dry). Dashed lines represent latitudinal zones with a day length variation of approximately 1.5 hours using the summer solstice (June 21st) as the reference day. (b) Categorization of the 869 sites in the final data subset used for the statistical analysis. (c) Distribution of rural (green) and urban (red) study sites, classified using the Global Human Settlement Layer (European Commission, 2023). Note that panels (a) and (b) do not use the same color patterns for bioclimatic regions.

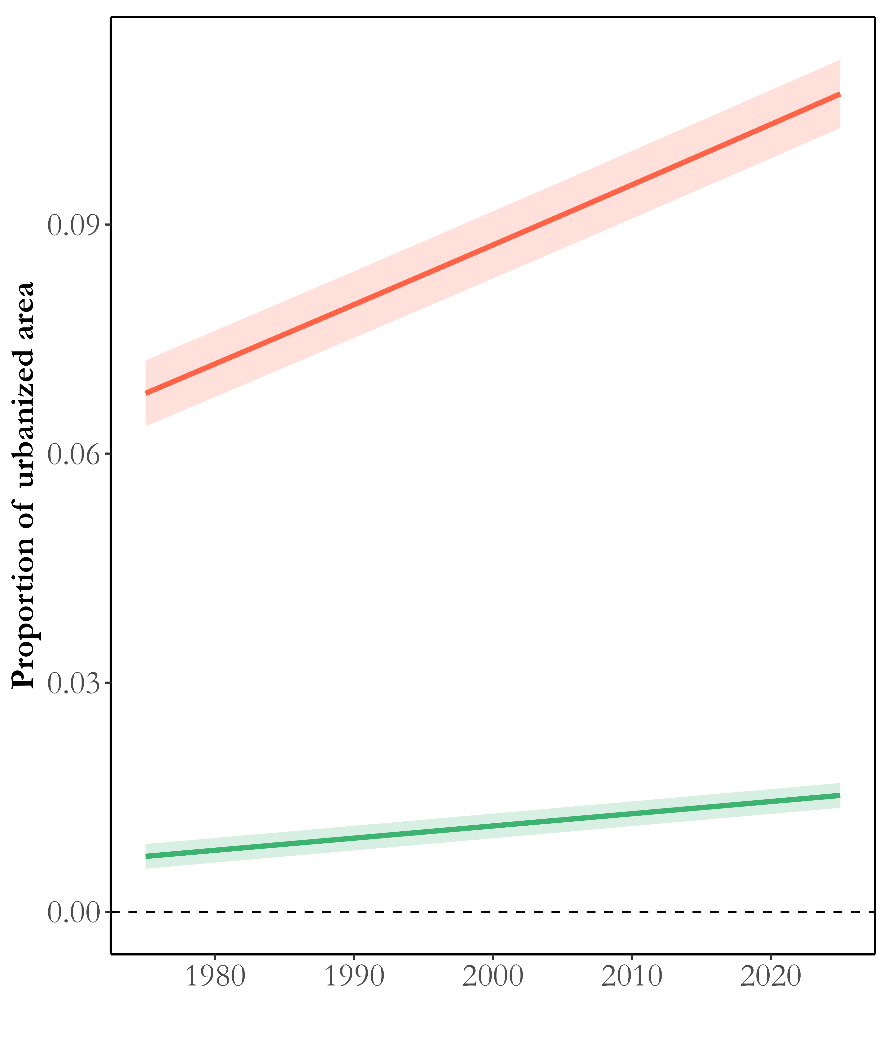

**Figure S2.** Temporal trends in the proportion of urbanized area for rural (green color) and urban sites (red color) from 1975 to 2025. The proportion of urbanized area was calculated as the percentage of urbanized area in 2x2 km grids. A linear mixed model was conducted with the proportion of urbanized area as the response variable, year as predictor variable and site as random factor. The shaded areas represent the 90% confidence intervals of the model predictions. This figure highlights the differences in the percentage of urbanized area between urban and rural sites and how these have changed over time.

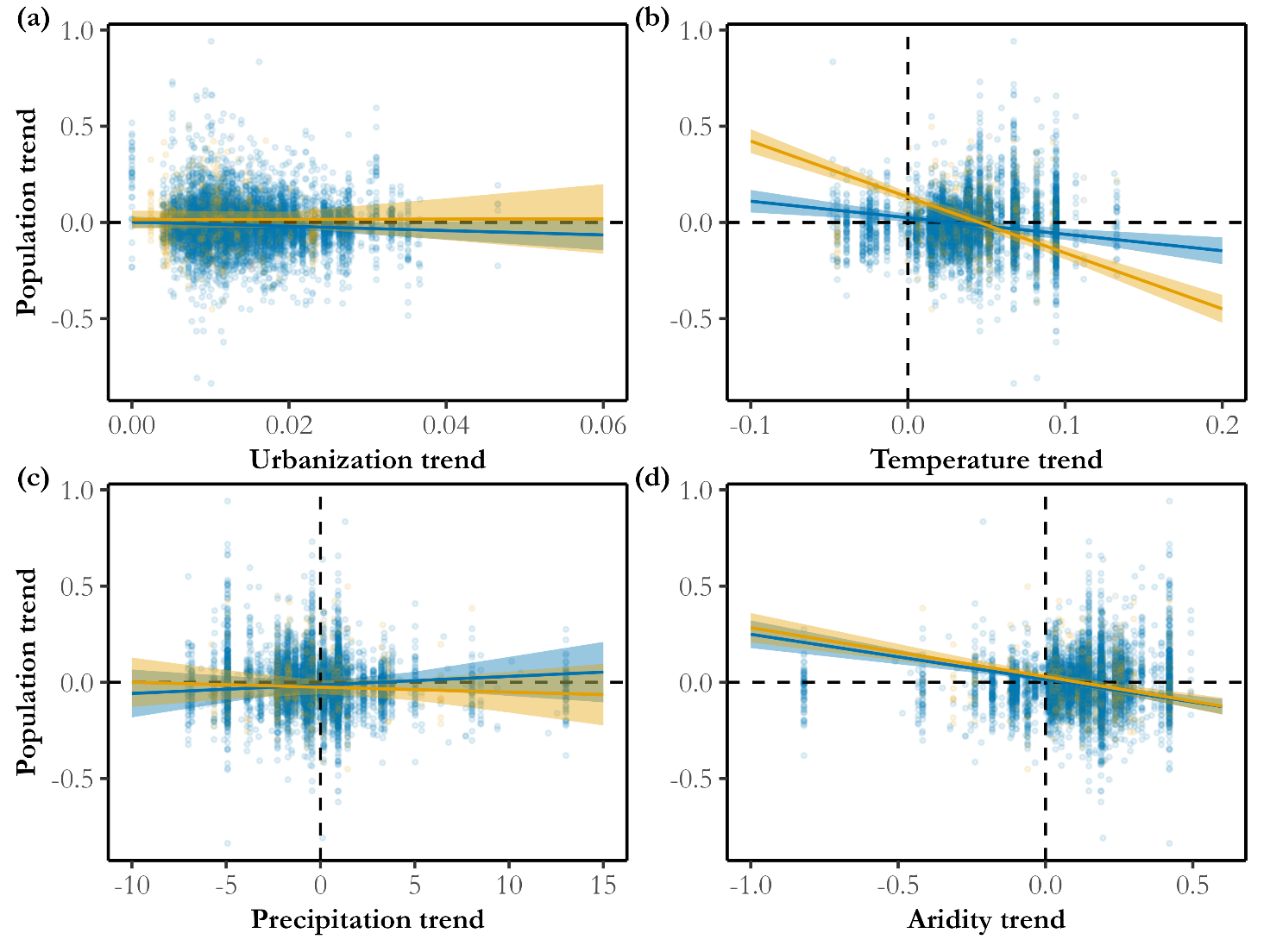

**Figure S3**. Relationships between butterfly population trends and urbanization and climate trends across rural (green) and urban (red) sites using all species available (i.e. not only common species in rural and urban sites: 145 species in rural sites and 100 species in urban sites). Points represent raw data, while lines indicate model predictions with 95% confidence intervals. The models tested the interactions between each predictor: (a) urbanization trend; (b) temperature trend; (c) precipitation trend; (d) aridity trend; and environmental type (rural vs urban). N = 8409 population trends.

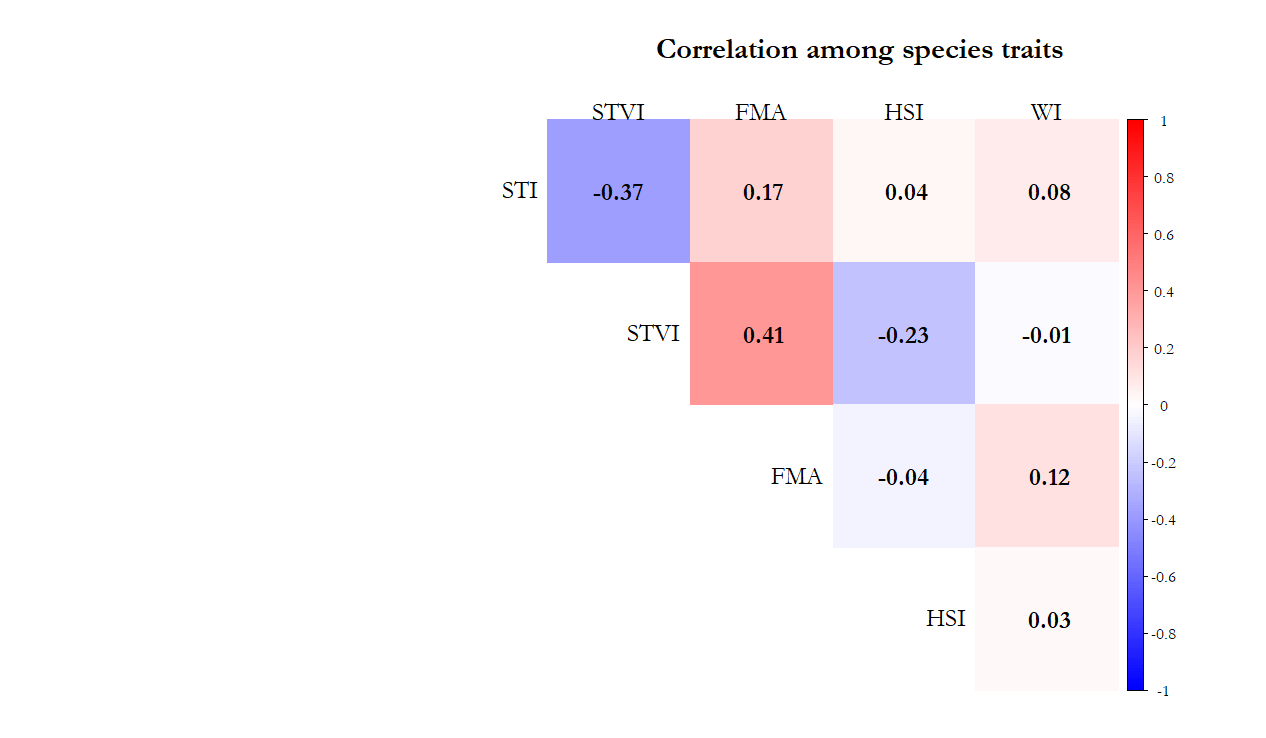

**Figure S4.** Correlation heatmap among species traits based on Pearson correlation coefficients. The plot shows the correlation values between five species traits: STI (species temperature index), STVI (species temperature variation index), FMA (flight month average), HSI (host-plant specialization index), and WI (wing index). Positive correlations are indicated in red, and negative correlations are shown in blue, with color intensity reflecting the strength of the correlation. Correlation values are displayed within each cell.

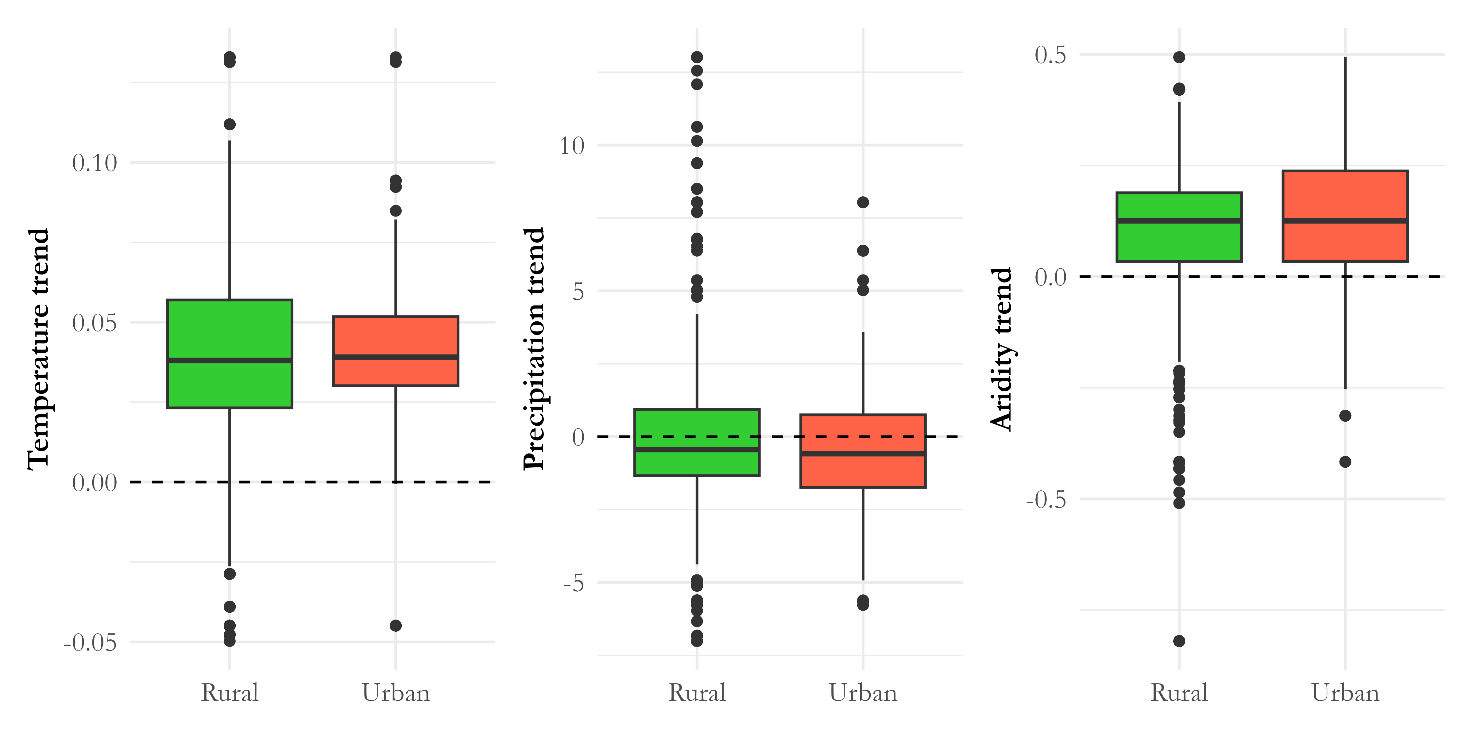

**Figure S5**. Boxplots showing the variation in (a) temperature trend, (b) precipitation trend, and (c) aridity trend across rural and urban sites. The dashed horizontal line at y = 0 indicates no trend. Rural sites are represented in green, and urban sites in red.

**SUPPLEMENTARY TABLES**

**TABLE S1**. Summary of the 145 butterfly species and their traits used in the analyses. Columns include the host plant index (HSI), wing index (WI), flight month average (FMA), species temperature index (STI), and species temperature variation index (STVI). The Urban column indicates whether each species is present in urban environments (at least one population following our criteria described in the methods). All species are present in rural sites. “NA” denotes missing data.

| **SPECIES** | **HSI** | **WI** | **FMA** | **STI** | **STVI** | **Urban** |
| --- | --- | --- | --- | --- | --- | --- |
| *Aglais io* | 0.236 | 0.061 | 7.5 | 8.84 | 2.91 | yes |
| *Aglais urticae* | 0.707 | 0.021 | 7 | 8.12 | 3.76 | yes |
| *Agriades optilete* | 0.218 | -0.046 | 3 | NA | NA | yes |
| *Anthocharis cardamines* | 0.101 | 0.001 | 4 | 8.3 | 3.88 | yes |
| *Anthocharis euphenoides* | 0.577 | -0.012 | 3.5 | 12.94 | 3.13 | no |
| *Apatura ilia* | 0.267 | 0.091 | 4.5 | 9.03 | 2.13 | no |
| *Aphantopus hyperantus* | 0.063 | 0.004 | 3 | 7.9 | 2.7 | yes |
| *Aporia crataegi* | 0.082 | 0.076 | 3.5 | 9.14 | 3.53 | yes |
| *Araschnia levana* | 0.707 | -0.009 | 5 | 8.62 | 1.97 | yes |
| *Arethusana arethusa* | 0.333 | 0.014 | 4 | 10.63 | 2.22 | no |
| *Argynnis pandora* | 0.577 | 0.109 | 5 | 11.92 | 3.06 | no |
| *Argynnis paphia* | 0.258 | 0.094 | 3.5 | 9.02 | 2.91 | yes |
| *Aricia agestis* | 0.102 | -0.048 | 7 | 10.16 | 2.89 | yes |
| *Aricia artaxerxes* | 0.056 | -0.049 | 4 | 6.45 | 4.12 | yes |
| *Aricia cramera* | 0.112 | -0.054 | 9.5 | 14.28 | 2.71 | yes |
| *Boloria dia* | 0.447 | -0.022 | 5.5 | 9.28 | 2.39 | yes |
| *Boloria euphrosyne* | 0.577 | -0.001 | 4 | 6.95 | 3.98 | yes |
| *Boloria selene* | 0.577 | -0.004 | 3.5 | 6.93 | 3.65 | yes |
| *Brenthis daphne* | 0.447 | 0.023 | 3.5 | 10.6 | 2.9 | yes |
| *Brenthis ino* | 0.204 | -0.008 | 2.5 | 6.86 | 3 | yes |
| *Brintesia circe* | 0.204 | 0.109 | 4.5 | 11.07 | 2.74 | yes |
| *Cacyreus marshalli* | 1 | -0.057 | 8 | 14.13 | 2.52 | yes |
| *Callophrys rubi* | 0.033 | -0.047 | 6 | 8.57 | 4.08 | yes |
| *Carcharodus alceae* | 0.167 | -0.041 | 8 | 11.14 | 3.23 | yes |
| *Carterocephalus palaemon* | 0.126 | -0.044 | 2.5 | 6.9 | 3.35 | yes |
| *Carterocephalus silvicola* | 0.2 | -0.055 | 2.5 | NA | NA | no |
| *Celastrina argiolus* | 0.02 | -0.039 | 7 | 9.14 | 3.81 | yes |
| *Charaxes jasius* | 0.102 | 0.148 | 6.5 | 14.77 | 1.96 | yes |
| *Coenonympha arcania* | 0.169 | -0.016 | 3.5 | 9.04 | 2.63 | yes |
| *Coenonympha dorus* | 0.169 | -0.027 | 3 | 12.67 | 2.96 | no |
| *Coenonympha glycerion* | 0.258 | -0.024 | 3 | 8.06 | 2.63 | yes |
| *Coenonympha pamphilus* | 0.096 | -0.041 | 7 | 8.96 | 3.89 | yes |
| *Coenonympha tullia* | 0.115 | -0.017 | 2 | 6.36 | 3.03 | no |
| *Colias alfacariensis* | 0.167 | 0.021 | 6.5 | 9.94 | 2.78 | yes |
| *Colias crocea* | 0.041 | 0.025 | 7 | 10.69 | 3.33 | yes |
| *Colias hyale* | 0.204 | 0.017 | 6.5 | 8.37 | 2.31 | yes |
| *Colias palaeno* | 0.707 | 0.029 | 3 | 3.62 | 3.25 | no |
| *Cupido alcetas* | 0.183 | -0.041 | 7 | 10.81 | 2.66 | no |
| *Cupido argiades* | 0.072 | -0.046 | 6 | 9.42 | 2.45 | yes |
| *Cupido minimus* | 0.105 | -0.062 | 6 | 8.76 | 3.26 | NA |
| *Cupido osiris* | 0.289 | -0.048 | 6 | 10.62 | 3 | no |
| *Cyaniris semiargus* | 0.169 | -0.038 | 6 | NA | NA | yes |
| *Erebia aethiops* | 0.043 | 0.022 | 3.5 | 8.1 | 2.25 | NA |
| *Erebia ligea* | 0.183 | 0.02 | 3 | 5.51 | 3.54 | yes |
| *Erebia medusa* | 0.224 | 0.002 | 3.5 | 8.4 | 2.27 | yes |
| *Erebia meolans* | 0.2 | 0.013 | 3.5 | 8.94 | 3.33 | no |
| *Erebia neoridas* | 0.289 | 0.002 | 2.5 | 10.06 | 2.91 | no |
| *Erynnis tages* | 0.129 | -0.042 | 4 | 9.12 | 2.9 | yes |
| *Euchloe crameri* | 0.154 | 0.008 | 3 | 12.79 | 3.13 | yes |
| *Eumedonia eumedon* | 0.218 | -0.037 | 4 | NA | NA | no |
| *Euphydryas aurinia* | 0.102 | -0.004 | 3.5 | 9.53 | 3.07 | yes |
| *Euphydryas maturna* | 0.046 | 0.002 | 2.5 | 7.64 | 2.7 | no |
| *Fabriciana adippe* | 0.5 | 0.051 | 3.5 | NA | NA | yes |
| *Fabriciana niobe* | 0.408 | 0.036 | 2.5 | NA | NA | yes |
| *Favonius quercus* | 0.408 | -0.032 | 5 | 9.49 | 2.79 | no |
| *Glaucopsyche alexis* | 0.065 | -0.036 | 5 | 9.59 | 3.59 | no |
| *Glaucopsyche melanops* | 0.224 | -0.046 | 2 | 13.26 | 2.86 | yes |
| *Gonepteryx cleopatra* | 0.577 | 0.061 | 7.5 | 13.95 | 2.66 | yes |
| *Gonepteryx rhamni* | 0.218 | 0.063 | 8 | 8.81 | 3.38 | yes |
| *Hamearis lucina* | 0.577 | -0.032 | 6 | 9.11 | 2.37 | yes |
| *Hesperia comma* | 0.333 | -0.036 | 3.5 | 8.47 | 3.19 | yes |
| *Heteropterus morpheus* | 0.25 | -0.025 | 3 | 9.52 | 2.18 | yes |
| *Hipparchia fagi* | 0.25 | 0.095 | 4.5 | 10.53 | 2.62 | no |
| *Hipparchia fidia* | 0.144 | 0.066 | 3.5 | 13.55 | 2.55 | yes |
| *Hipparchia hermione* | 0.408 | 0.074 | 2 | 10.34 | 2.92 | yes |
| *Hipparchia semele* | 0.143 | 0.036 | 3.5 | 9.18 | 2.61 | yes |
| *Hipparchia statilinus* | 0.123 | 0.03 | 5 | 11.83 | 2.99 | yes |
| *Iphiclides feisthamelii* | 0.408 | 0.12 | 7.5 | NA | NA | yes |
| *Iphiclides podalirius* | 0.105 | 0.118 | 6 | 10.87 | 3.4 | yes |
| *Issoria lathonia* | 0.408 | 0.011 | 7 | 9.33 | 3.08 | yes |
| *Lampides boeticus* | 0.058 | -0.03 | 9 | 12.82 | 3.03 | yes |
| *Lasiommata maera* | 0.072 | 0.027 | 6 | 8.56 | 3.76 | yes |
| *Lasiommata megera* | 0.12 | 0.013 | 6.5 | 10.39 | 3.13 | yes |
| *Lasiommata petropolitana* | 0.258 | 0.001 | 3.5 | 5.07 | 3.19 | no |
| *Leptidea juvernica* | NA | 0.008 | 3 | NA | NA | no |
| *Leptidea reali* | 1 | 0 | 6 | NA | NA | no |
| *Leptidea sinapis* | 0.144 | -0.003 | 6 | 9.11 | 3.92 | yes |
| *Leptotes pirithous* | 0.045 | -0.053 | 8 | 12.82 | 3.37 | yes |
| *Libythea celtis* | 0.707 | 0.006 | 6.5 | 12.19 | 2.91 | no |
| *Limenitis camilla* | 0.289 | 0.064 | 3 | 8.85 | 2.26 | no |
| *Limenitis reducta* | 0.577 | 0.037 | 5 | 11.07 | 3.26 | yes |
| *Lycaena hippothoe* | 0.289 | -0.027 | 4.5 | 6.45 | 3.4 | yes |
| *Lycaena phlaeas* | 0.333 | -0.048 | 9.5 | 9.29 | 3.94 | yes |
| *Lycaena tityrus* | 0.136 | -0.036 | 7 | 9.35 | 2.59 | yes |
| *Lycaena virgaureae* | 0.408 | -0.035 | 5 | 7.27 | 3.01 | yes |
| *Lysandra bellargus* | 0.183 | -0.031 | 7 | NA | NA | yes |
| *Lysandra coridon* | 1 | -0.026 | 3 | NA | NA | yes |
| *Lysandra hispana* | 1 | -0.023 | 7 | NA | NA | no |
| *Maniola jurtina* | 0.289 | 0.022 | 5 | 9.85 | 3.29 | yes |
| *Melanargia galathea* | 0.107 | 0.031 | 3.5 | 9.71 | 2.55 | no |
| *Melanargia lachesis* | 0.136 | 0.041 | 2.5 | 13.09 | 2.25 | yes |
| *Melanargia occitanica* | 0.154 | 0.039 | 2.5 | 13.79 | 2.26 | no |
| *Melitaea athalia* | 0.107 | -0.004 | 3.5 | 8.27 | 3.55 | yes |
| *Melitaea cinxia* | 0.289 | -0.003 | 4 | 9.6 | 2.89 | yes |
| *Melitaea deione* | 0.12 | -0.004 | 4.5 | 11.58 | 3.96 | no |
| *Melitaea didyma* | 0.051 | -0.005 | 5.5 | 10.42 | 3.1 | yes |
| *Melitaea parthenoides* | 0.105 | -0.019 | 3 | 10.61 | 2.66 | no |
| *Melitaea phoebe* | 0.154 | 0.013 | 4.5 | 10.91 | 3.19 | yes |
| *Melitaea trivia* | 0.447 | -0.017 | 5 | 10.97 | 2.71 | no |
| *Minois dryas* | 0.2 | 0.066 | 3 | 9.52 | 2.31 | no |
| *Muschampia proto* | 0.577 | -0.033 | 4 | NA | NA | no |
| *Nymphalis antiopa* | 0.192 | 0.106 | 5.5 | 7.61 | 3.69 | yes |
| *Nymphalis polychloros* | 0.136 | 0.069 | 6.5 | 9.68 | 3 | yes |
| *Ochlodes sylvanus* | 0.083 | -0.035 | 4 | 8.58 | 3.31 | yes |
| *Papilio machaon* | 0.053 | 0.116 | 6.5 | 9.28 | 4.08 | yes |
| *Pararge aegeria* | 0.154 | 0.008 | 7 | 9.71 | 3.44 | yes |
| *Phengaris nausithous* | 1 | -0.015 | 3 | 8.39 | 1.37 | yes |
| *Phengaris teleius* | 1 | -0.021 | 3 | 8.6 | 1.92 | yes |
| *Pieris brassicae* | 0.115 | 0.067 | 7 | 9.29 | 3.83 | yes |
| *Pieris mannii* | 0.408 | 0.018 | 7.5 | 11.46 | 3.13 | yes |
| *Pieris napi* | 0.114 | 0.015 | 5.5 | 8.21 | 4 | yes |
| *Pieris rapae* | 0.118 | 0.02 | 9.5 | 9.63 | 3.67 | yes |
| *Plebejus argus* | 0.069 | -0.049 | 4 | 8.61 | 3.48 | yes |
| *Plebejus argyrognomon* | 0.224 | -0.034 | 4 | 9.51 | 2.23 | yes |
| *Plebejus idas* | 0.041 | -0.046 | 5 | 6.68 | 4.07 | yes |
| *Polygonia c-album* | 0.05 | 0.029 | 6 | NA | NA | yes |
| *Polyommatus amandus* | 0.204 | -0.022 | 3 | 7.66 | 3.35 | yes |
| *Polyommatus celina* | 1 | -0.04 | 11 | NA | NA | no |
| *Polyommatus escheri* | 0.5 | -0.029 | 3 | 10.87 | 3.38 | no |
| *Polyommatus fulgens* | 1 | -0.032 | 2 | 11.1 | 0.93 | no |
| *Polyommatus icarus* | 0.144 | -0.035 | 7 | 9.07 | 4.11 | yes |
| *Polyommatus ripartii* | 0.447 | -0.033 | 4 | 10.96 | 2.53 | no |
| *Polyommatus thersites* | 0.5 | -0.04 | 7 | 10.59 | 2.99 | no |
| *Pontia daplidice* | 0.061 | 0.007 | 5.5 | 10.43 | 3.55 | yes |
| *Pontia edusa* | 0.102 | 0.011 | 5.5 | NA | NA | yes |
| *Pseudophilotes panoptes* | 0.5 | -0.066 | 6 | NA | NA | no |
| *Pyrgus armoricanus* | 0.183 | -0.049 | 5.5 | 10.7 | 2.97 | no |
| *Pyrgus malvae* | 0.192 | -0.058 | 5 | 8.74 | 3.37 | yes |
| *Pyrgus malvoides* | NA | -0.061 | 3.5 | NA | NA | no |
| *Pyronia bathseba* | 0.354 | -0.002 | 4.5 | 13.54 | 2.84 | yes |
| *Pyronia cecilia* | 0.177 | -0.024 | 3.5 | 14.06 | 2.48 | yes |
| *Pyronia tithonus* | 0.081 | -0.007 | 3.5 | 10.86 | 2.64 | yes |
| *Satyrium acaciae* | 1 | -0.036 | 3 | 10.21 | 2.67 | no |
| *Satyrium esculi* | 0.707 | -0.029 | 4 | 13.33 | 2.95 | yes |
| *Satyrium ilicis* | 0.112 | -0.025 | 4 | 10.21 | 2.91 | no |
| *Satyrium pruni* | 0.134 | -0.033 | 3 | 8.31 | 2.27 | yes |
| *Satyrium spini* | 0.167 | -0.033 | 3.5 | 10.22 | 3.21 | no |
| *Speyeria aglaja* | 0.5 | 0.057 | 3.5 | NA | NA | yes |
| *Spialia sertorius* | 1 | -0.059 | 7 | 10.44 | 3.36 | no |
| *Thymelicus acteon* | 0.118 | -0.056 | 4.5 | 11.31 | 3.23 | yes |
| *Thymelicus lineola* | 0.067 | -0.052 | 3.5 | 8.69 | 3.19 | yes |
| *Thymelicus sylvestris* | 0.101 | -0.045 | 4.5 | 9.87 | 2.96 | yes |
| *Vanessa atalanta* | 0.316 | 0.065 | 8.5 | 9.07 | 3.85 | yes |
| *Vanessa cardui* | 0.083 | 0.055 | 8 | 9.04 | 4.12 | yes |
| *Zerynthia rumina* | 0.577 | 0.025 | 5 | 13.87 | 2.83 | no |

**TABLE S2**. Complete results of the GLMMs testing interaction effects of environmental trends and site type (rural vs. urban) on butterfly population trends. Test statistics for models ordered based on AIC are shown: coefficient estimate (Coef), standard error (SD), z-value (Z), p-value (p), variation in Akaike's Information Criterion (ΔAIC), conditional R^2^ and marginal R^2^. All models have a sample size of 7,731 observations, 100 species, and 846 sites corresponding to 6 different climate regions.

| **Model predictor** | **Coef[SD]** | **Z(p)** | **ΔAIC** | **R^2^_c_** | **R^2^_m_** |
| --- | --- | --- | --- | --- | --- |
| Urbanization | -1.35[1.06] | -1.27(0.2) | 15026 | 0.842 | 0.007 |
| Site type | 0.01[0.03] | 0.37(0.71) |  |  |  |
| Urbanization * Site type | 1.05[2.14] | 0.49(0.62) |  |  |  |
| Temperature | -1.14[0.01] | -3.53(<0.01) | 0 | 0.886 | 0.06 |
| Site type | 0.09[0.01] | 7.92(<0.01) |  |  |  |
| Temperature * Site type | -1.73[0.06] | -27.4(<0.01) |  |  |  |
| Precipitation | 0.002[<0.01] | 0.24(0.81) | 5983 | 0.975 | 0.001 |
| Site type | -0.02[0.02] | -0.54(0.59) |  |  |  |
| Precipitation * Site type | -0.01[<0.01] | -15.6(<0.01) |  |  |  |
| Aridity | -0.26[0.05] | -4.8(<0.01) | 21059 | 0.902 | 0.052 |
| Site type | 0.01[0.012] | 0.68(0.49) |  |  |  |
| Aridity * Site type | 0.03[0.01] | 2.53(0.01) |  |  |  |

**TABLE S3**. Results of GLMMs testing the effect of urbanization on rural populations and urban populations, separately. For these models we used all available species in each data subset: 100 species in urban sites and 145 species in rural sites.

| **Environment type** | **Coefficient** | **Standard Error** | **Z-value** | **p-value** |
| --- | --- | --- | --- | --- |
| Rural | -1.11 | 1.09 | 1.02 | 0.31 |
| Urban | -4.21 | 5.31 | -0.79 | 0.428 |

**TABLE S4**. Complete results of the GLMMs testing interaction effects of environmental trends and species traits on butterfly population trends. Each interaction was tested separately for rural and urban sites. All model variables were standardized to a mean of 0 and a standard deviation of 1, allowing us to compare interaction effect sizes between each trend (urbanization, temperature, precipitation and aridity) and the different species traits. Test statistics for each model term are shown: Coefficient estimate (Coef), standard error (SD), z-value (Z), p-value(p), Akaike's information criterion (AIC), variation in AIC (ΔAIC), conditional R^2^ and marginal R^2^. Models for rural sites have a sample size of 6328 observations, 126 species and 688 sites corresponding to 6 different climate regions. Models for urban sites have a sample size of 841 observations, 90 species and 108 sites corresponding to 5 different climate regions.

| **Data**  **subset** | **Model predictor** | **Coef [SD]** | **Z (p)** | **AIC** | **ΔAIC** | **R^2^_c_** | **R^2^_m_** |
| --- | --- | --- | --- | --- | --- | --- | --- |
| Rural  sites | Urbanization | -0.0032 [0.0043] | -0.738 (0.46) | -15227574 | 1518 | 0.787 | 0.001 |
|  | HSI | -0.0004 [0.0026] | -0.172 (0.86) |  |  |  |  |
|  | Urbanization * HSI | 0.0004 [0.0003] | 13.78 (<0.01) |  |  |  |  |
|  | Urbanization | -0.0031 [0.0043] | -0.727 (0.467) | -15227594 | 1498 | 0.787 | 0.001 |
|  | WI | 0.0029 [0.0033] | 0.89(0.371) |  |  |  |  |
|  | Urbanization * WI | 0.0004 [0.0003] | 14.47 (<0.01) |  |  |  |  |
|  | Urbanization | -0.0028 [0.0043] | -0.67 (0.5) | -15228453 | 639 | 0.787 | 0.011 |
|  | FMA | 0.0126 [0.0034] | 3.73 (<0.01) |  |  |  |  |
|  | Urbanization * FMA | 0.0009 [<0.0001] | 32.49 (<0.01) |  |  |  |  |
|  | Urbanization | -0.0031 [0.0042] | -0.76 (0.44) | -15227509 | 1583 | 0.787 | 0.002 |
|  | STI | 0.004 [0.0034] | 1.58 (0.11) |  |  |  |  |
|  | Urbanization * STI | 0.0003[<0.0001] | -11.1 (<0.01) |  |  |  |  |
|  | Urbanization | -0.0031 [0.0043] | -1.46 (0.14) | -15229092 | 0 | 0.787 | 0.001 |
|  | STVI | 0.0017 [0.003] | 0.56 (0.58) |  |  |  |  |
|  | Urbanization * STVI | 0.0018[<0.001] | 41.3 (<0.01) |  |  |  |  |
|  | Temperature | -0.0302 [0.0006] | -51.50 (<0.01) | -14592062 | 653 | 0.809 | 0.057 |
|  | HSI | -0.0008 [0.0025] | -0.34 (0.73) |  |  |  |  |
|  | Temperature * HSI | -0.0018 [<0.0001] | -39.92 (<0.01) |  |  |  |  |
|  | Temperature | -0.0304 [0.0006] | -51.86 (<0.01) | -14591162 | 1510 | 0.809 | 0.058 |
|  | WI | 0.0034 [0.0033] | 1.03 (0.3) |  |  |  |  |
|  | Temperature * WI | 0.0011 [<0.0001] | 27.13 (<0.01) |  |  |  |  |
|  | Temperature | -0.0282 [0.0006] | -48.02 (<0.01) | -14591443 | 1142 | 0.809 | 0.06 |
|  | FMA | 0.0129 [0.0034] | 3.81 (<0.01) |  |  |  |  |
|  | Temperature * FMA | 0.0014 [0.0000] | 33.04 (<0.01) |  |  |  |  |
|  | Temperature | -0.0287 [0.0006] | -48.99 (<0.01) | -14592624 | 0 | 0.808 | 0.054 |
|  | STI | 0.0046 [0.0027] | 1.70 (0.09) |  |  |  |  |
|  | Temperature * STI | 0.0023 [0.0001] | 47.39 (<0.01) |  |  |  |  |
|  | Temperature | -0.0286 [0.0006] | -48.65 (<0.01) | -14591803 | 895 | 0.807 | 0.051 |
|  | STVI | 0.0026 [0.0031] | 0.82 (0.41) |  |  |  |  |
|  | Temperature * STVI | 0.0015 [<0.0001] | 36.77 (<0.01) |  |  |  |  |
|  | Precipitation | 0.0262 [0.0004] | 60.67 (<0.01) | -14598760 | 854 | 0.81 | 0.043 |
|  | HSI | -0.0002 [0.0025] | -0.08 (0.94) |  |  |  |  |
|  | Precipitation * HSI | -0.0035 [<0.0001] | -82.74 (<0.01) |  |  |  |  |
|  | Precipitation | 0.0279 [0.0004] | 64.46 (<0.01) | -14592012 | 7595 | 0.813 | 0.048 |
|  | WI | 0.0029 [0.0033] | 0.89 (0.37) |  |  |  |  |
|  | Precipitation * WI | -0.0004 [<0.0001] | -10.04 (<0.01) |  |  |  |  |
|  | Precipitation | 0.0276 [0.0004] | 63.82 (<0.01) | -14592085 | 6531 | 0.813 | 0.055 |
|  | FMA | 0.0126 [0.0034] | 3.74 (<0.01) |  |  |  |  |
|  | Precipitation * FMA | -0.0005 [<0.0001] | -12.33 (<0.01) |  |  |  |  |
|  | Precipitation | 0.029 [0.0004] | 67.12 (<0.01) | -14599635 | 0 | 0.816 | 0.051 |
|  | STI | 0.0039 [0.0027] | 1.42 (0.16) |  |  |  |  |
|  | Precipitation * STI | -0.0039 [<0.0001] | -87.75 (<0.01) |  |  |  |  |
|  | Precipitation | 0.029 [0.0004] | 67.01 (<0.01) | -14597566 | 2032 | 0.815 | 0.05 |
|  | STVI | 0.0022 [0.0031] | 0.70 (0.48) |  |  |  |  |
|  | Precipitation * STVI | 0.0031 [<0.0000] | 75.28 (<0.01) |  |  |  |  |
|  | Aridity | -0.0362 [0.0005] | -74.62 (<0.01) | -14596135 | 7866 | 0.822 | 0.076 |
|  | HSI | -0.0001 [0.0025] | -0.06 (0.95) |  |  |  |  |
|  | Aridity * HSI | 0.0022 [<0.0000] | 49.78 (<0.01) |  |  |  |  |
|  | Aridity | -0.0378 [0.0005] | -77.77 (<0.01) | -14594367 | 9636 | 0.824 | 0.082 |
|  | WI | 0.0030 [0.0033] | 0.93 (0.35) |  |  |  |  |
|  | Aridity * WI | 0.0011 [<0.0000] | 26.59 (<0.01) |  |  |  |  |
|  | Aridity | -0.0367 [0.0005] | -75.56 (<0.01) | -14594555 | 9421 | 0.823 | 0.086 |
|  | FMA | 0.0127 [0.0034] | 3.78 (<0.01) |  |  |  |  |
|  | Aridity * FMA | 0.0014 [0.0001] | 30.15 (<0.01) |  |  |  |  |
|  | Aridity | -0.0385 [0.0005] | -79.29 (<0.01) | -14603967 | 0 | 0.827 | 0.083 |
|  | STI | 0.0046 [0.0027] | 1.69 (0.09) |  |  |  |  |
|  | Aridity * STI | 0.0049 [0.0001] | 101.75 (<0.01) |  |  |  |  |
|  | Aridity | -0.0380 [0.0005] | 78.3 (<0.01) | -14595728 | 8280 | 0.825 | 0.082 |
|  | STVI | 0.0021 [0.0031] | 0.68 (0.49) |  |  |  |  |
|  | Aridity * STVI | -0.002 [<0.0000] | -45.42 (<0.01) |  |  |  |  |
| Urban  sites | Urbanization | -0.0005 [0.0096] | 0.05 (0.958) | -1767429 | 2813 | 0.855 | 0.006 |
|  | HSI | -0.0099 [0.0066] | -1.5 (0.132) |  |  |  |  |
|  | Urbanization * HSI | -0.003[<0.0001] | 34.26 (<0.01) |  |  |  |  |
|  | Urbanization | 0.0006 [0.0097] | 0.07 (0.945) | -1770242 | 0 | 0.855 | 0.002 |
|  | WI | 0.0026 [0.0077] | 0.34 (0.732) |  |  |  |  |
|  | Urbanization * WI | -0.0051 [<0.0001] | -63.3 (<0.01) |  |  |  |  |
|  | Urbanization | 0.0018 [0.0097] | 0.19 (0.845) | -1768563 | 1678 | 0.855 | 0.003 |
|  | FMA | 0.0048 [0.0084] | 0.57 (0.566) |  |  |  |  |
|  | Urbanization * FMA | 0.0045 [<0.0001] | 48.1 (<0.01) |  |  |  |  |
|  | Urbanization | 0.0015 [0.0096] | 0.165 (0.869) | -1766269 | 3973 | 0.853 | 0.003 |
|  | STI | 0.0063 [0.0056] | 1.12 (0.26) |  |  |  |  |
|  | Urbanization * STI | -0.0003 [<0.0001] | -3.65 (<0.01) |  |  |  |  |
|  | Urbanization | 0.0018 [0.0097] | 0.189 (0.85) | -1766938 | 3304 | 0.855 | 0.004 |
|  | STVI | 0.0077 [0.0074] | 1.03 (0.3) |  |  |  |  |
|  | Urbanization * STVI | 0.0024 [<0.0001] | 26.13 (<0.01) |  |  |  |  |
|  | Temperature | -0.0608 [0.0013] | -45.59 (<0.01) | -1766612 | 1489 | 0.896 | 0.157 |
|  | HSI | -0.0087 [0.0066] | -1.32 (0.19) |  |  |  |  |
|  | Temperature * HSI | 0.0009 [0.0001] | 8.11 (<0.01) |  |  |  |  |
|  | Temperature | -0.0613 [0.0013] | -45.94 (<0.01) | -1766547 | 1554 | 0.897 | 0.154 |
|  | WI | 0.0002 [0.0078] | 0.03 (0.98) |  |  |  |  |
|  | Temperature * WI | -0.0002 [<0.0001] | -1.50 (0.13) |  |  |  |  |
|  | Temperature | -0.0595 [0.0013] | -44.45 (<0.01) | -1766748 | 1353 | 0.894 | 0.15 |
|  | FMA | 0.0073 [0.0085] | 0.85 (0.39) |  |  |  |  |
|  | Temperature * FMA | 0.0017 [<0.0001] | 14.24 (<0.01) |  |  |  |  |
|  | Temperature | -0.0568 [0.0013] | -42.49 (<0.01) | -1768102 | 0 | 0.892 | 0.142 |
|  | STI | 0.0074 [0.0057] | 1.29 (0.19) |  |  |  |  |
|  | Temperature * STI | 0.0051 [<0.0001] | 39.48 (<0.01) |  |  |  |  |
|  | Temperature | -0.0628 [0.0013] | -46.97 (<0.01) | -1766837 | 1265 | 0.898 | 0.165 |
|  | STVI | 0.0069 [0.0075] | 0.92 (0.36) |  |  |  |  |
|  | Temperature * STVI | -0.0023 [<0.0001] | -17.06 (<0.01) |  |  |  |  |
|  | Precipitation | 0.0056 [0.0010] | 5.46 (<0.01) | -1771927 | 0 | 0.861 | 0.015 |
|  | HSI | -0.0112 [0.0068] | -1.64 (0.101) |  |  |  |  |
|  | Precipitation * HSI | 0.0090 [<0.0001] | 86.20 (<0.01) |  |  |  |  |
|  | Precipitation | 0.0114 [0.0010] | 11.08 (<0.01) | -1764617 | 7309 | 0.852 | 0.007 |
|  | WI | 0.0004 [0.0079] | 0.06 (0.95) |  |  |  |  |
|  | Precipitation * WI | -0.0009 [<0.0001] | -8.59 (<0.01) |  |  |  |  |
|  | Precipitation | 0.0059 [0.0010] | 5.76 (<0.01) | -1766483 | 5442 | 0.851 | 0.007 |
|  | FMA | 0.0076 [0.0086] | 0.88 (0.38) |  |  |  |  |
|  | Precipitation * FMA | 0.0045 [<0.0001] | 44.08 (<0.01) |  |  |  |  |
|  | Precipitation | 0.0106 [0.0010] | 10.33 (<0.01) | -1764545 | 7381 | 0.851 | 0.008 |
|  | STI | 0.0058 [0.0058] | 0.99 (0.32) |  |  |  |  |
|  | Precipitation * STI | -0.0001 [<0.0001] | -0.77 (0.44) |  |  |  |  |
|  | Precipitation | 0.0115 [0.0010] | 11.29 (<0.01) | -1765738 | 6187 | 0.852 | 0.012 |
|  | STVI | 0.0067 [0.0075] | 0.90 (0.37) |  |  |  |  |
|  | Precipitation * STVI | 0.0041 [<0.0001] | 34.58 (<0.01) |  |  |  |  |
|  | Aridity | -0.0336 [0.0013] | -26.13 (<0.01) | -1769827 | 0 | 0.878 | 0.066 |
|  | HSI | -0.0111 [0.0068] | -1.64 (0.1) |  |  |  |  |
|  | Aridity * HSI | -0.0075 [<0.0001] | -67.08 (<0.01) |  |  |  |  |
|  | Aridity | -0.0400 [0.0013] | -30.87 (<0.01) | -1765431 | 4395 | 0.877 | 0.078 |
|  | WI | 0.0004 [0.0079] | 0.06 (0.96) |  |  |  |  |
|  | Aridity * WI | 0.0010 [<0.0001] | 9.39 (<0.01) |  |  |  |  |
|  | Aridity | -0.0354 [0.0013] | -27.28 (<0.01) | -1765903 | 3923 | 0.872 | 0.067 |
|  | FMA | 0.0071 [0.0086] | 0.82 (0.41) |  |  |  |  |
|  | Aridity * FMA | -0.0024 [<0.0001] | -23.66 (<0.01) |  |  |  |  |
|  | Aridity | -0.0388 [0.0013] | -30.06 (<0.01) | -1765642 | 4184 | 0.875 | 0.074 |
|  | STI | 0.0059 [0.0058] | 1.02 (0.31) |  |  |  |  |
|  | Aridity * STI | 0.0019 [<0.0001] | 17.27 (<0.01) |  |  |  |  |
|  | Aridity | -0.0408 [<0.0013] | -31.62 (<0.01) | -1766420 | 3407 | 0.877 | 0.085 |
|  | STVI | 0.0067 [0.0075] | 0.90 (0.37) |  |  |  |  |
|  | Aridity * STVI | -0.0039 [<0.0001] | -32.80 (<0.01) |  |  |  |  |
